# Integrated Multi-Omics Analysis for Molecular Subtyping in NSCLC: A Cohort Study

**DOI:** 10.1101/2025.09.05.674586

**Authors:** Bin Yan, Zhangyu Wang, Yun Peng, Kefu Liu, Jian-Guo Zhou, Shixiang Wang

**Affiliations:** Department of Biomedical Informatics, School of Life Sciences, Central South University, Changsha, China; Department of Urology, Peking University People’s hospital, Beijing, China; Department of Oncology, The Second Affiliated Hospital of Zunyi Medical University, Zunyi, China

**Author notes:** Correspondence (J.-G.Z.); (S.W.).

**Keywords:** non-small cell lung cancer, molecular subtyping, multi-omics integration, unsupervised clustering

## Abstract

Lung cancer remains the leading cause of cancer-related morbidity and mortality worldwide, with non-small cell lung cancer (NSCLC) accounting for approximately 85% of cases. Current histopathological classification and driver gene testing provide limited prognostic and therapeutic guidance due to intra-tumoral heterogeneity and incomplete characterization of the tumor microenvironment (TME). Here, we constructed single- and multi-omics molecular classification systems for NSCLC by integrating transcriptomic, genomic, epigenomic, proteomic, and TME data from TCGA LUAD and LUSC cohorts. Single-omics analyses revealed distinct molecular patterns, including DNA methylation subtypes associated with sex and histology. Multi-omics integration identified five consensus subtypes closely corresponding to histology, with three immune-activated (CS2–CS4) and two immunosuppressive (CS1, CS5) subtypes. While overall survival did not differ significantly, progression-free survival analysis highlighted CS4 as a high-risk subtype with *KRAS*-driven genomic alterations. These findings provide a framework for NSCLC molecular stratification and highlight molecular and immune features that could guide future research on targeted therapies and immunotherapy.

## 1. Introduction

Lung cancer is one of the most prevalent malignancies worldwide. According to the GLOBOCAN 2022 data released by International Agency for Research on Cancer, in 2022, lung cancer had the highest incidence and estimated mortality rates among all cancers, making it once again the world’s leading cancer[1]. Non-small cell lung cancer (NSCLC), which accounts for approximately 85% of lung cancer cases[2], has an overall 5-year survival rate of less than 20%. While early screening can reduce mortality, it is associated with a high false-positive rate. Moreover, the postoperative recurrence rate for stage I patients can reach 20%–40%, and the 5-year survival rate after recurrence and metastasis is only approximately 15%[3]. These phenomena highlight the limitations of current diagnostic and therapeutic approaches.

Currently, the clinical diagnosis and treatment of NSCLC rely primarily on histopathological classification and the Tumor-Node-Metastasis (TNM) staging system. However, this classification system has some limitations. For example, patients with morphologically identical lung squamous cell carcinoma may have completely different prognoses due to distinct driver gene mutations (e.g., *NFE2L2, ASAH2*)[4]. Some early-stage patients may still experience recurrence due to molecular abnormalities, such as *EGFR* mutations, despite having a low tumor stage[5]. Conversely, some late-stage patients may benefit from targeted or immunotherapy due to specific molecular characteristics[6].

While driver gene testing has spurred progress in precision medicine for NSCLC, it is limited by several inherent constraints. These include: (1) a narrow testing scope, which predominantly focuses on a few well-established targets such as *EGFR, ALK*, and *ROS1*[7], potentially overlooking the broader molecular landscape of tumors; (2) the absence of identifiable driver mutations in a subset of patients, thereby excluding them from benefiting from targeted therapies; and (3) the near-inevitability of acquired resistance, exemplified by the development of resistance to osimertinib—a third-generation *EGFR* tyrosine kinase inhibitor (TKI)—within 2–3 years in most patients[8–10]. Furthermore, the information provided by driver gene testing alone is insufficient to reflect the complex interplay of factors such as the tumor microenvironment (TME), epigenetic modifications, and transcriptomic profiles, which collectively influence prognosis and treatment responses. Consequently, the traditional diagnostic and treatment model, which relies predominantly on histopathological classification, TNM staging, and driver gene testing, falls short of meeting the precise classification and treatment demands for NSCLC.

In recent years, multi-omics integration research has emerged as a new approach, covering genomics, transcriptomics, epigenomics, and proteomics, and offering a more comprehensive perspective for characterizing the molecular heterogeneity of NSCLC[11,12]. However, while some studies have classified tumors using TME as an independent omics layer[13–15], there is still a lack of systematic classification attempts that incorporate TME into a multi-omics integration system. Notably, key components of the TME, including immune cell infiltration and angiogenesis, are strongly associated with the survival outcomes of NSCLC patients and their responsiveness to immunotherapy, and may further modulate the efficacy of targeted therapies[16,17].

Based on these considerations, our study integrates multi-omics data from The Cancer Genome Atlas (TCGA) lung adenocarcinoma (LUAD) and lung squamous cell carcinoma (LUSC) cohorts, including transcriptomics, somatic mutations, copy number variations, DNA methylation, and Reverse Phase Protein Array (RPPA) protein expression. In addition, we independently derived TME features from bulk transcriptome data. By performing consensus clustering at the single-omics level and applying ten multi-omics integration algorithms to construct consensus subtypes, we aim to establish a more comprehensive NSCLC classification system. This framework is intended to provide novel insights for precision diagnosis and treatment by systematically assessing the clinical prognosis and biological characteristics of different subtypes.

## 2. Results

### 2.1. Identification of NSCLC Molecular Subtypes by Single-Omics Consensus Clustering

To identify NSCLC molecular subtypes, we first examined six types of omics data, revealing pronounced differences in gene expression, DNA methylation, copy number variation, and protein expression, as generally expected for these histological subtypes (Supplementary Materials, Figure S1).

Unsupervised consensus clustering of gene expression data classified NSCLC samples into four transcriptomic subtypes (Figure 1). And the optimal number of clusters was determined using the cumulative distribution function (CDF), Delta Area Plot, and Tracking Plot.. Clustering results for the other omics data indicated that, except for CNV data which favored five clusters, all other omics data were best represented by four clusters (Supplementary Materials, Figures S2–S6). These single-omics subtypes exhibited pronounced omics-specific features, providing a foundation for subsequent multi-omics integrative analysis.

**Figure 1.**
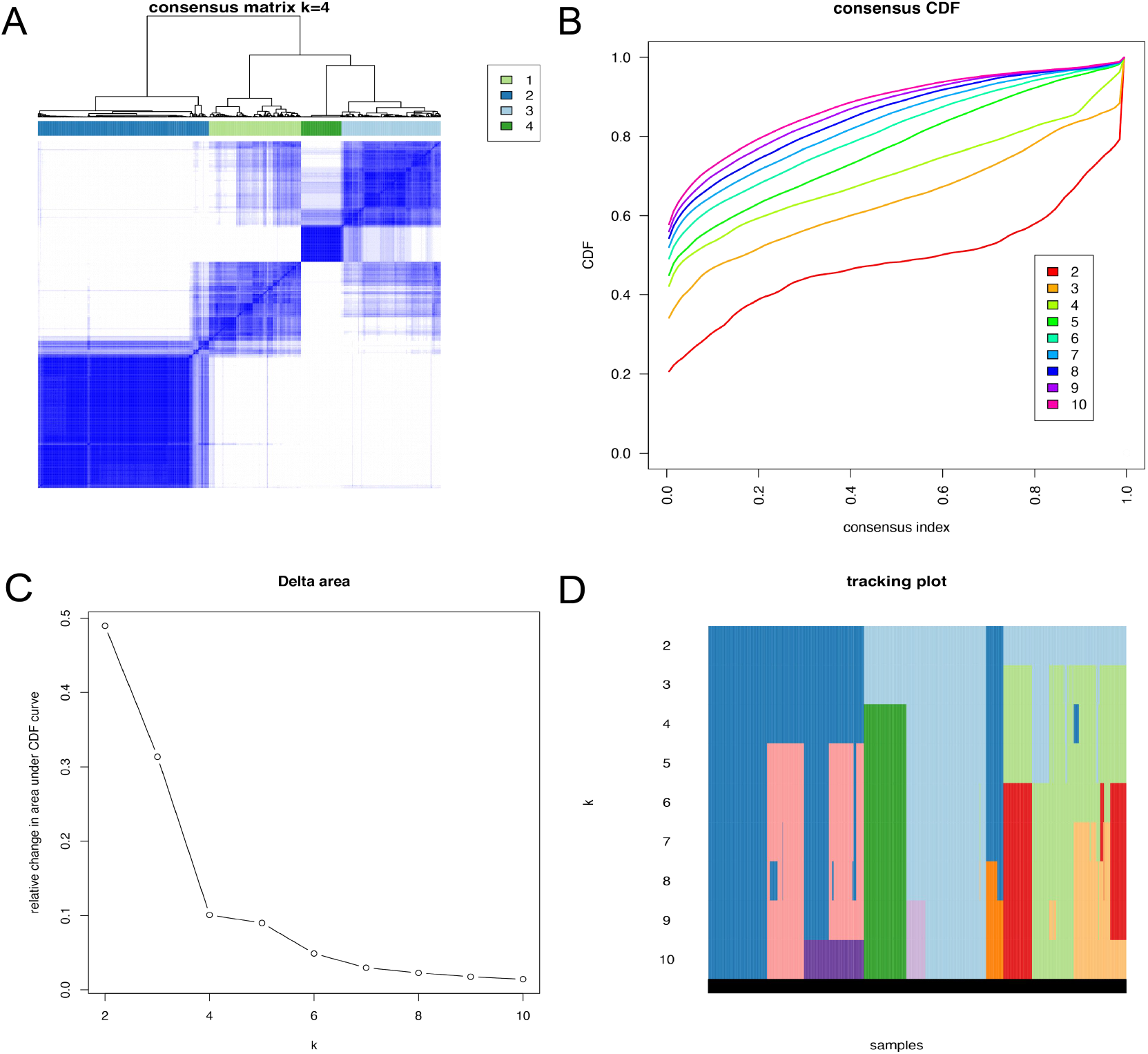
Consensus clustering results of gene expression data. (**A**) Consensus matrix heatmap at K = 4; (**B**) Cumulative distribution function plot; (**C**) Relative change in the area under the CDF curve (Delta Area Plot); (**D**) Sample classification tracking plot across different values of K.

### 2.2. Integrative Multi-Omics Clustering Reveals Five Robust NSCLC Subtypes

The optimal number of clusters for the multi-omics dataset was determined by calculating the clustering prediction index (CPI) and Gap statistic, both of which reached relatively high values at K = 5, indicating the best model performance at this resolution (Supplementary Materials, Figure S7).

Following this, consensus clustering was performed using ten machine learning algorithms. A total of 551 NSCLC samples were stratified into five robust multi-omics consensus subtypes (CS1–CS5), with the following sample sizes and proportions: CS1, 116 cases (21.1%); CS2, 113 cases (20.5%); CS3, 140 cases (25.4%); CS4, 125 cases (22.7%); and CS5, 57 cases (10.3%) (Figure 2). Clustering quality assessed by silhouette coefficients (Figure 2C) showed that CS2 and CS4 exhibited higher average widths, indicating that these clusters represent well-defined molecular subgroups suitable for downstream analyses. In contrast, CS3 displayed a relatively low silhouette width (0.25), suggesting lower within-cluster similarity and potential overlap with neighboring subtypes.

**Figure 2.**
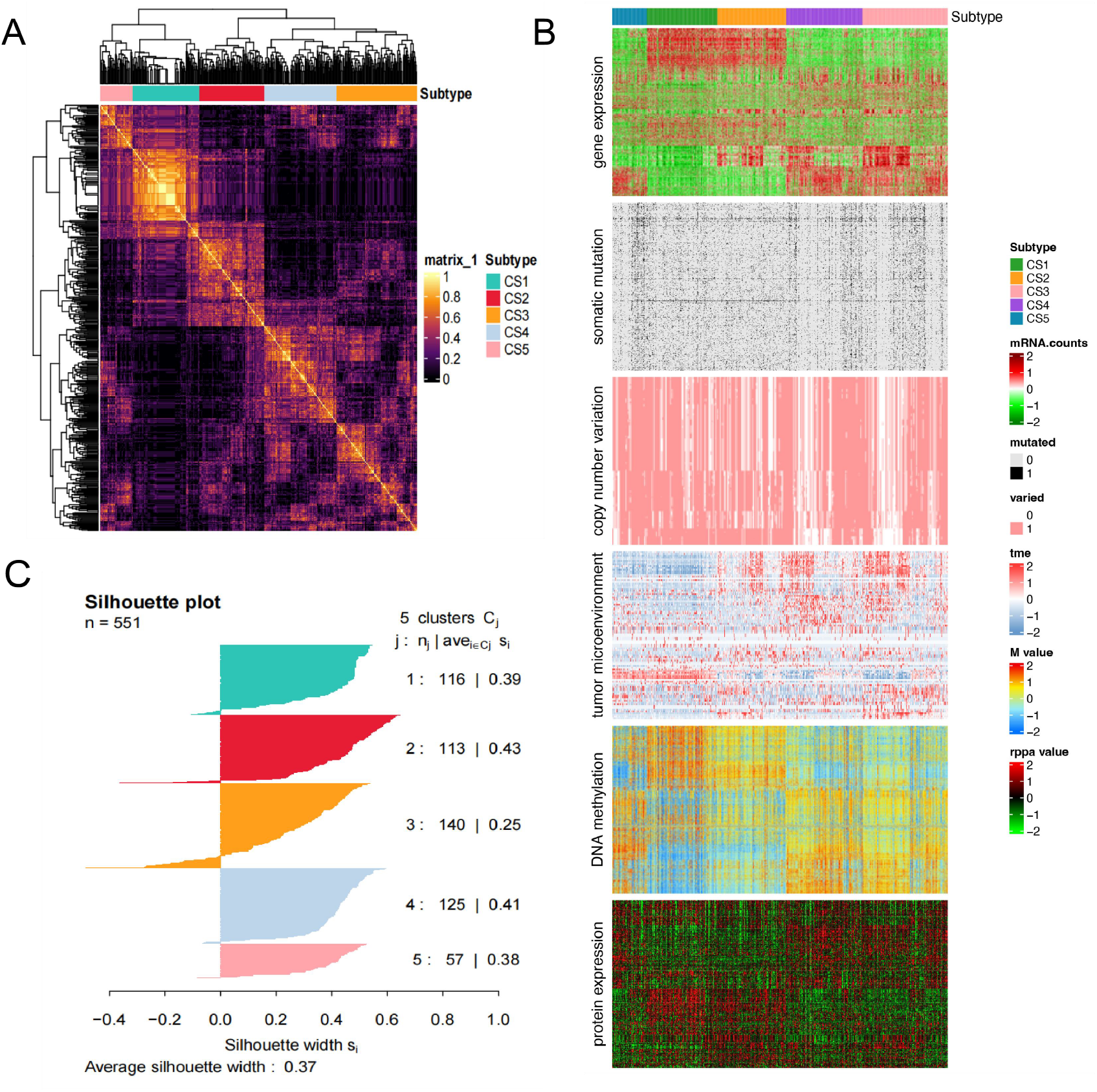
Multi-omics consensus subtypes of NSCLC. (**A**) Consensus matrix heatmap of the five subtypes; (**B**) Molecular landscapes of the five subtypes across six omics layers; (**C**) Clustering quality assessment based on silhouette coefficients.

### 2.3. Cross-Omics Consistency and Clinical Relevance of Single- and Multi-Omics Subtypes in NSCLC

To explore the associations between multi-omics consensus subtypes, single-omics subtypes, and clinical features (including histological type, clinical stage, and gender), we performed heatmap visualization (Figure 3A). The results revealed that certain subtype classifications (e.g., MOVICS, CNV, and MET) were strongly associated with histological type. Notably, the MET subtypes also showed strong correlations with patient gender, whereas proteomic and tumor microenvironment subtypes exhibited weaker associations.

**Figure 3.**
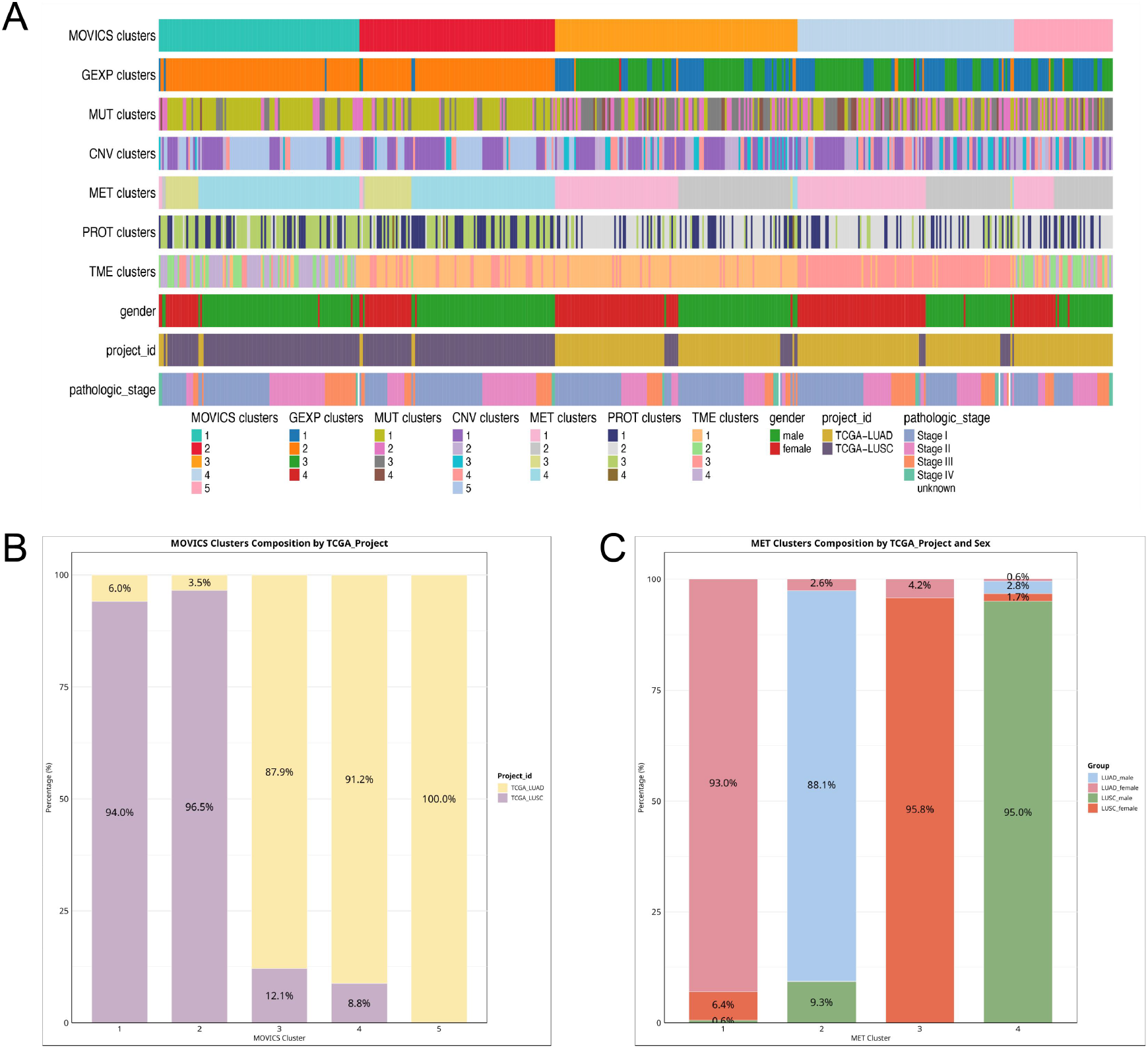
Associations of MOVICS and single-omics subtypes with clinical features. (**A**) Heatmap showing the concordance between MOVICS consensus subtypes, single-omics clustering results, and clinical features (gender, project, and stage); (**B**) Distribution of histological subtypes (LUAD vs. LUSC) across the MOVICS consensus subtypes. (**C**) Association of DNA methylation (MET) subtypes with histological type and sex.

We further examined the composition of histological types across different subtype results using stacked bar plots (Figure 3B, 3C, and Supplementary Figure S8). The multi-omics classification divided NSCLC into five consensus subtypes, which were highly correlated with histological type (Figure 3B). LUAD samples predominantly clustered in MOVICS clusters 3, 4, and 5, comprising 108 (87.9%), 114 (91.2%), and 57 (100%) patients, respectively. LUSC samples were mainly distributed in clusters 1 and 2, with 109 (94.0%) and 109 (96.5%) patients, respectively. These five subtypes were subsequently designated CS1–CS5 for downstream analyses.

For single-omics subtypes (excluding DNA methylation), GEXP, CNV, MUT, PROT, and TME clusters exhibited diverse enrichment patterns: some clusters were LUAD-dominant (e.g., GEXP 1,3; CNV 2–4), others LUSC-dominant (e.g., GEXP 2; CNV 5; MUT 1), and several displayed a relatively balanced distribution between the two histological types (e.g., PROT 1; TME 1,2) (Supplementary Materials, Figure S8).

Figure 3C further shows the distribution of the four methylation subtypes stratified by histological type and gender. MET clusters 1 and 2 were enriched in LUAD, while clusters 3 and 4 were enriched in LUSC. Methylation subtypes displayed striking associations with both histology and gender: MET cluster 1 comprised mostly female LUAD (93.0%), cluster 2 was predominantly male LUAD (88.1%), cluster 3 was mostly female LUSC (95.8%), and cluster 4 was predominantly male LUSC (95.0%).

### 2.4. Prognostic Impact of Multi-Omics Subtypes on Survival Outcomes

In the overall survival (OS) analysis, the Kaplan–Meier curves showed no significant differences among the five multi-omics consensus subtypes defined by MOVICS (Log-rank *p* = 0.53) (Figure 4A). This lack of separation may reflect the limited sample size or insufficient biological heterogeneity across subtypes, resulting in reduced statistical power. In the multivariable Cox regression analysis, after adjusting for clinical stage, clinical stage emerged as an independent prognostic factor for OS. In contrast, the hazard ratios of the consensus subtypes were modest, and the corresponding confidence intervals encompassed 1, suggesting no independent prognostic value (Figure 4C).

**Figure 4.**
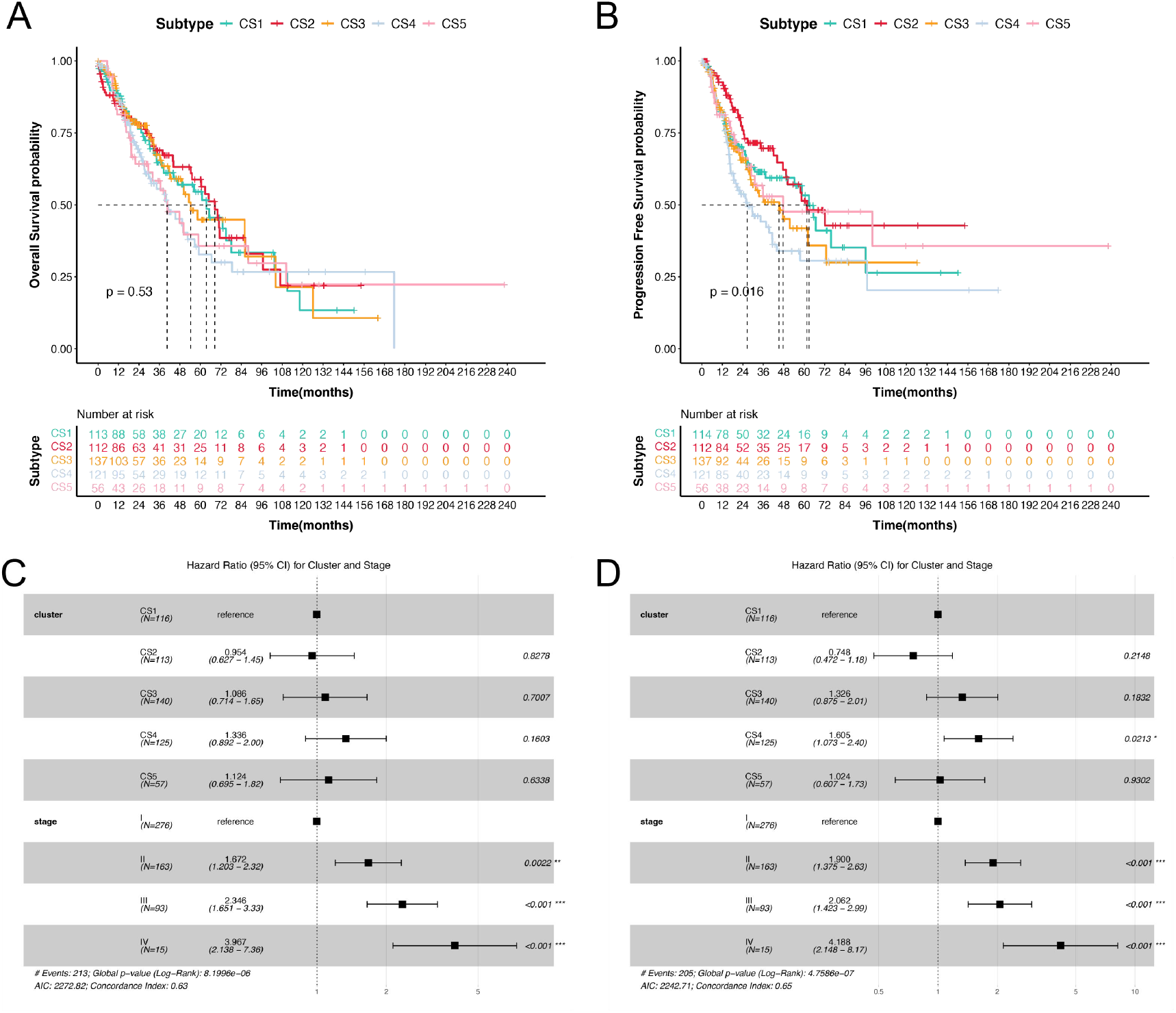
Survival analysis of multi-omics consensus subtypes: Kaplan-Meier curves and Cox regression results for OS and PFS. (**A**) Kaplan–Meier curves of OS stratified by multi-omics consensus subtypes; (**B**) Kaplan-Meier curves of PFS stratified by multi-omics consensus subtypes; (**C)** Multivariable Cox regression analysis for OS (consensus subtypes and clinical stage); (**D**) Multivariable Cox regression analysis for PFS (consensus subtypes and clinical stage).

Notably, the progression-free survival (PFS) analysis revealed significant differences among the five subtypes (Log-rank *p*= 0.016) (Figure 4B). Multivariable Cox regression revealed that the CS4 subtype had the highest risk (HR = 1.605, 95% CI 1.075–2.40, *p* = 0.0213), although its confidence interval was relatively wide. Compared with CS1, the hazard ratios for CS2, CS3, and CS5 were not statistically significant (Figure 4D).

### 2.5. Subtype-Specific Molecular Characteristics and Functional Pathway Landscapes

#### 2.5.1. Mutational Landscape Reveals Subtype-specific Alterations

Genetic mutations play a pivotal role in tumor initiation and progression[18]. In this study, we compared the mutational landscapes of the five multi-omics consensus subtypes. As shown in Figure 5A, a total of 18 significantly mutated genes were identified across the five consensus subtypes. *TP53, TTN, SYNE1, FAM135B, KMT2D*, and *LRRK2* were more frequently mutated in CS1 and CS2, which were predominantly composed of LUSC, whereas *KRAS, ASXL3, STK11, EGFR, LPPR4*, and *CLCN1* were more commonly mutated in CS3–CS5, which were enriched for LUAD. Supplementary Table 1 further highlights subtype-specific mutational profiles: CS1 was enriched for *TTN, FAM135B, NPAP1*, and *CDKN2A* mutations; CS2 for *TP53, SYNE1, KMT2D, LRRK2*, and *NFE2L2*; CS4 for *KRAS* and *EGFR*; and CS5 for *KEAP1, ASXL3, STK11*, and *CLCN1*.

**Figure 5.**
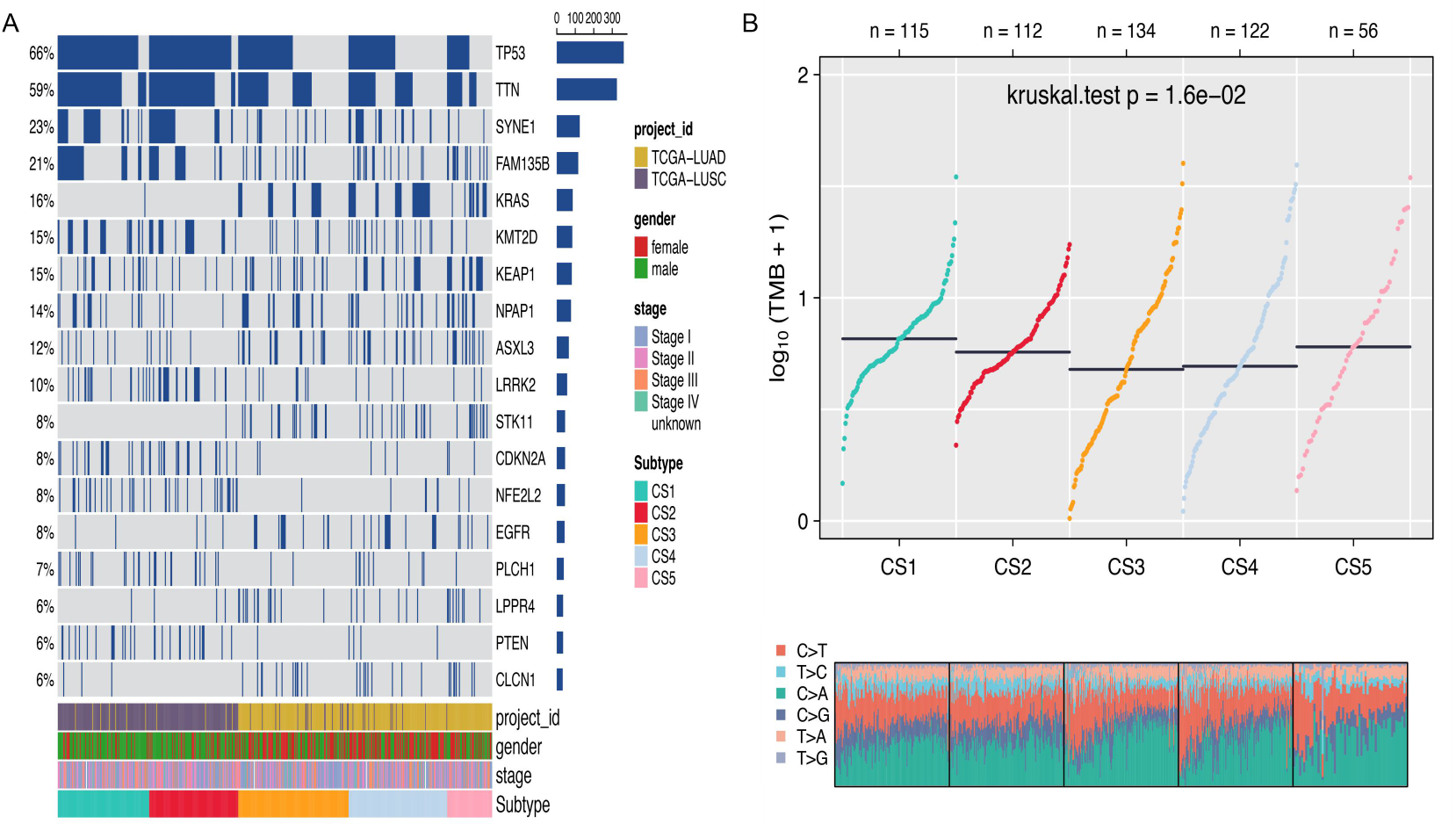
Mutation landscape of the five multi-omics consensus subtypes. (A) Waterfall plots of significantly mutated genes in each subtype; (B) Comparison of TMB and Ti/Tv ratios across subtypes.

Tumor mutational burden (TMB), a surrogate for neoantigen load, has been linked to improved response to immune checkpoint inhibitors[19,20]. The transition-to-transversion (Ti/Tv) ratio also carries clinical significance in NSCLC; for instance, smoking-related C>A transversions are associated with poor prognosis, whereas *APOBEC*-driven C>T transitions may predict immunotherapy sensitivity[21]. In our analysis, TMB differed significantly across subtypes (p < 0.05), with CS1 highest and CS3 lowest. LUSC-dominated subtypes (CS1, CS2) had higher TMB than LUAD-dominated ones (CS3–CS5), suggesting greater genomic instability. Ti/Tv analysis further showed a higher proportion of T>C transitions in CS1 and CS2.

#### 2.5.2. Distinct Molecular Signatures of Multi-Omics Subtypes

Through differential expression analysis, we identified subtype-specific genes across the five multi-omics consensus subtypes (Supplementary Materials, Figure S9A). Members of the keratin gene family (e.g., *KRT5, KRT6A, KRT6B*) were markedly upregulated in the LUSC-dominant subtypes (CS1 and CS2) but downregulated in the LUAD-dominant subtypes (CS3 – CS5). From these analyses, we selected 200 subtype-specific signature genes for each subtype (100 upregulated and 100 downregulated). Their distinct expression profiles across subtypes supported the robustness of our classification and provided a molecular basis for external validation (Supplementary Materials, Figure S9B, S9C).

#### 2.5.3. Subtype-Specific Functional Pathways

We performed subtype-specific gene set enrichment analyses (GSEA) for each of the five multi-omics subtypes and then compared the enrichment patterns across subtypes. KEGG pathway analysis (Figure 6) revealed that CS2, CS3, and CS4 consistently exhibited upregulation in pathways including “Th17 cell differentiation,” “Chemokine signaling pathway,” “Cytokine–cytokine receptor interaction,” and “Phagosome,” whereas CS1 and CS5 showed consistent downregulation, suggesting the presence of two distinct immune-related subtype groups. GO term enrichment comparison (Supplementary Materials, Figure S10) further demonstrated that gene sets associated with “leukocyte-mediated immunity,” “immunoglobulin production,” “lymphocyte-mediated immunity,” and “B cell-mediated immunity” were consistently upregulated in CS2 and CS3 and downregulated in CS1 and CS5, largely corroborating the KEGG enrichment patterns.

**Figure 6.**
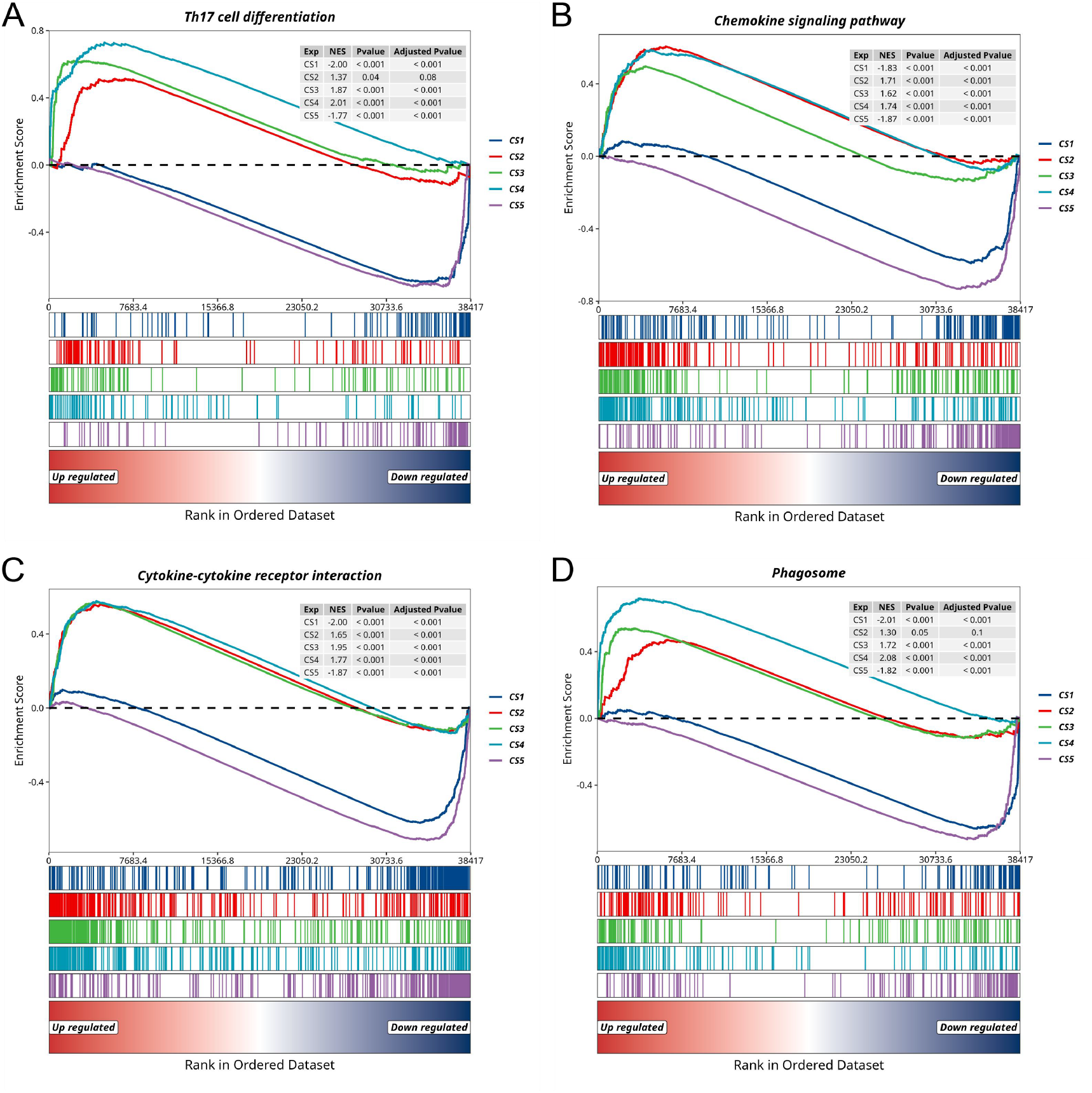
Comparison of selected immune-related KEGG pathway enrichment among five multi-omics consensus subtypes. (**A**) KEGG pathway enrichment for “Th17 cell differentiation”; (**B**) KEGG pathway enrichment for “Chemokine signaling pathway”; (**C**) KEGG pathway enrichment for “Cytokine–cytokine receptor interaction” ;(**D**) KEGG pathway enrichment for “Phagosome”.

## 3. Discussion

In this study, we constructed single- and multi-omics molecular classification systems for NSCLC by integrating transcriptomic, genomic, epigenomic, proteomic, and tumor microenvironment data. Analyses at the single-omics level revealed distinct patterns: DNA methylation subtypes were strongly associated with sex and histology, likely reflecting smoking-induced methylation alterations[22], whereas proteomic subtypes were largely independent of histopathological classification. This suggests that tumors of different origins may converge on similar functional modules, including proliferation, immune evasion, and metabolic adaptation.

Integration of these multi-omics layers yielded consensus subtypes that corresponded closely with histology: CS1 and CS2 were predominantly LUSC, whereas CS3–CS5 were mainly LUAD. Beyond tissue-of-origin characteristics, the subtypes exhibited consistent immune profiles across histological subtypes. Specifically, CS2, CS3, and CS4 showed immune-activated features, whereas CS1 and CS5 displayed immunosuppressive traits, suggesting the presence of two distinct immune subtypes that may differ in their responses to immunotherapy.

Despite these pronounced molecular distinctions, OS did not differ significantly among subtypes. This lack of association may reflect several factors, including the high proportion of early-stage patients (I–II, 79.5%), unaccounted treatment heterogeneity, and the limited sample size of CS5 (n = 57). Interestingly, PFS analysis revealed that CS4 was associated with higher risk, consistent with its *KRAS*-driven genomic alterations. These findings suggest that patients in this subtype could potentially benefit from MEK inhibitor-based combination immunotherapy[23].

Nevertheless, several limitations warrant consideration: (1) The TCGA cohort includes only LUAD and LUSC, excluding rare subtypes such as large-cell neuroendocrine carcinoma and sarcomatoid carcinoma, which harbor distinctive molecular alterations (e.g., TP53 and *STK11/KEAP1* co-mutations or *RB1/TP53* co-deletion)[24]. Their absence may limit the biological generalizability of the classification. (2) The lack of microbiome and metabolome data may constrain the completeness of subtype characterization[25]. (3) Key fusion events (*ALK, ROS1, RET*) were not systematically incorporated; for example, *CD74-ROS1* rearrangements are common in *ROS1*-positive LUAD and predict sensitivity to crizotinib[26], and their omission may result in misclassification of fusion-driven subtypes.(4) TCGA lacks targeted therapy and immune checkpoint inhibitor response data, preventing evaluation of subtype-treatment associations and limiting clinical translational potential.

Importantly, some planned analyses were not completed in the present study. Future work will aim to: (1) optimize single-omics clustering to identify more robust and biologically meaningful subtypes; (2) systematically compare multi-omics subtypes with established TCGA classifications; (3) validate the classification in independent external cohorts; and (4) further characterize the genomic and tumor microenvironment features of the multi-omics consensus subtypes. These efforts are expected to refine NSCLC molecular stratification and provide a more reliable framework for precision oncology.

## 4. Materials and Methods

### 4.1. Data Source and Preprocessing

We downloaded TCGA non-small cell lung cancer (NSCLC) data from the UCSC Xena website (https://xena.ucsc.edu/), including LUAD and LUSC cohorts: RNA-seq gene expression data (counts format, n = 1129, normalized and log2-transformed across pan-cancer samples); DNA methylation data (Illumina Human Methylation 450K, 503 LUAD samples and 412 LUSC samples); copy number variation (CNV) data (Gene Level Copy Number, ASCAT3, 503 LUAD samples and 490 LUSC samples); protein expression data (RPPA, 365 LUAD samples and 328 LUSC samples); and somatic mutation data (Pan-Cancer Atlas, 513 LUAD samples and 480 LUSC samples, binary encoding with 1 indicating nonsynonymous mutations and 0 indicating wild type).

To reduce data dimensionality and facilitate clustering analysis, the preprocessing and feature selection of the other five omics datasets were performed as follows: (1) Somatic mutation data: retain genes mutated in at least 5% of samples; (2) Protein expression data: remove proteins with missing values; (3) RNA-seq data: Z-score normalization for each gene across samples, followed by selection of highly variable genes based on median absolute deviation (MAD); (4) CNV data: rank genes based on the variation frequency, defined as the proportion of samples with copy number not equal to 2 (derived from ASCAT3); (5) DNA methylation data: rank probes based on standard deviation to select highly variable loci.

In addition, clinical phenotypes, survival data, and TPM expression data were downloaded from UCSC Xena for downstream analyses, while MAF-formatted mutation data were obtained separately using the TCGAblinks R package[27]. For simplicity in downstream analyses and figure annotations, the six omics layers are referred to by the following variable names: GEXP (gene expression), MUT (somatic mutation), CNV (copy number variation), MET (DNA methylation), PROT (protein expression), and TME (tumor microenvironment features).

### 4.2. TME Feature Calculation

TME features were computed based on RNA-seq data downloaded from the GDC portal (https://portal.gdc.cancer.gov/) for TCGA LUAD and LUSC cohorts. Seven built-in immune infiltration algorithms implemented in the IOBR R package (CIBERSORT, ESTIMATE, quanTIseq, TIMER, IPS, MCPCounter, and EPIC) were applied to quantify the infiltration levels of various immune cell types and other TME characteristics for each sample[28]. These features were subsequently incorporated into the multi-omics integration analysis and used for subtype construction.

### 4.3. Identification of Single-Omics Molecular Subtypes

Unsupervised clustering of the six omics datasets was performed using the ConsensusClusterPlus R package with hierarchical clustering[29]. The number of features used for clustering in each omics dataset was as follows: the top 20% highly variable genes for RNA-seq (4,106), the top 1% most variable probes for DNA methylation (8,064), the top 10% genes ranked by variation frequency for CNV (5,794), 651 somatic mutation features, 206 protein expression features, and 96 TME features.

Clustering parameters were set as follows: maxK = 10, reps = 1000, pItem = 0.8, pFeature = 1, clusterAlg = “hc”, innerLinkage = “ward.D2”, and finalLinkage = “ward.D2”. The distance parameter was adjusted according to the omics type: “binary” for mutation data, “euclidean” for CNV data, and “pearson” for all other omics types.

### 4.4. Multi-Omics Consensus Clustering

A total of 551 samples with available data across six omics layers were stratified using the MOVICS package for multi-omics consensus clustering[30]. Missing values were imputed and high-variance features were selected using the built-in “getElites” function. The parameters of “getElites*”* were configured to balance the number of features across the different omics layers: the top 3,000 features with the highest MAD were retained for RNA-seq and DNA methylation data, 50% of proteins with the highest MAD (243 proteins) were retained for RPPA data, all 96 TME features were retained, and the top 3,000 CNV features with the highest standard deviation were selected. CNV values were binarized such that samples with copy number not equal to 2 were considered as having a CNV, and those equal to 2 as no CNV. For somatic mutation data, 619 genes mutated in at least 5% of samples were retained.

The optimal number of clusters (K) was determined based on the CPI and gap statistics. Subsequently, the ten multi-omics integration algorithms implemented in the MOVICS package (intNMF, moCluster, SNF, PINSPlus, NEMO, CIMLR, COCA, iClusterBayes, ConsensusClustering, and LRAcluster) were applied to derive clustering solutions, followed by consensus integration across these methods to obtain robust molecular subtypes. Subtype stability was evaluated using silhouette scores.

### 4.5. Evaluation of Genetic Alterations among Different Subtypes

The “compMut” function from the MOVICS package was applied to analyze mutation features among subtypes, identifying significantly different and frequently mutated genes (mutation frequency >5%, p < 0.05). TMB was calculated as the number of nonsynonymous mutations per million bases.

### 4.6. Subtype-Specific Transcriptomic Profiling

Differential expression analysis among subtypes was performed using the “runDEA” function from the MOVICS package, with the limma method selected for statistical testing. Multi-omics volcano plots were generated using the jjVolcano function from the scRNAtoolVis package[31]. Subtype-specific biomarkers were further identified by selecting genes with the most significant log2 fold changes (adjusted p < 0.05) while ensuring that the selected biomarkers were not shared with other subtypes.

### 4.7. Gene Set Enrichment Analysis

Differentially expressed genes were ranked according to log2 fold change (log2FC). GSEA was conducted with the R package clusterProfiler for GO and KEGG pathways[32,33]. Enrichment results were visualized using enrichPlot and GseaVis to characterize the biological functions and signaling pathways associated with each subtype[34].

### 4.8. Statistical Analysis

All data preprocessing, statistical analyses, and visualizations were performed in the R environment (version 4.4.1). Kaplan–Meier survival curves and Cox proportional hazards regression were performed using the survival package, and forest plots were generated with survminer. All tests were two-sided, with p < 0.05 considered statistically significant.

## Supplementary Materials

The following supporting information can be downloaded at: https://www.mdpi.com/article/doi/s1, Figure S1: title; Table S1: title; Video S1: title.

## Author Contributions

Author Contributions: Conceptualization, S.W. and J.-G.Z.; methodology, B.Y., Z.W.; investigation, B.Y.; formal analysis, B.Y.; visualization, B.Y., Z.W. and Y.P.; writing— original draft preparation, B.Y.; writing— review and editing, B.Y., Z.W., Y.P., K.L., S.W. and J.-G.Z.; supervision, S.W. and J.-G.Z.; project administration, S.W. and K.L.; funding acquisition, S.W. and J.-G.Z. All authors have read and agreed to the published version of the manuscript.

## Institutional Review Board Statement

Not applicable. Ethical review and approval were waived for this study, as all data were obtained from the publicly available TCGA database, for which patient consent and ethical approval were already secured by the original investigators.

## Informed Consent Statement

Informed Consent Statement: Not applicable.

## Data Availability Statement

Publicly available datasets were analyzed in this study. TCGA data are available from The Cancer Genome Atlas (https://portal.gdc.cancer.gov/), and processed TCGA datasets were accessed via the UCSC Xena platform (https://xena.ucsc.edu/). No new data were generated in this study.

## Acknowledgments

We are grateful for resources from the Bioinformatics Platform, Furong Laboratory and Bioinformatics Center, Xiangya Hospital, Central South University.

## Abbreviations

The following abbreviations are used in this manuscript:

CNV: Copy number variation
CPI: Clustering prediction index
CS: Consensus subtype
GSEA: Gene set enrichment analysis
LUAD: Lung adenocarcinoma
LUSC: Lung squamous cell carcinoma
MAD: Median absolute deviation
NSCLC: Non-small cell lung cancer
OS: Overall survival
PFS: Progression-free survival
RPPA: Reverse phase protein array
TCGA: The Cancer Genome Atlas
Ti/Tv: The transition-to-transversion
TKI: Tyrosine kinase inhibitor
TMB: Tumor mutational burden
TME: Tumor microenvironment
TNM: Tumor-Node-Metastasis

## Disclaimer/Publisher’s Note

The statements, opinions and data contained in all publications are solely those of the individual author(s) and contributor(s) and not of MDPI and/or the editor(s). MDPI and/or the editor(s) disclaim responsibility for any injury to people or property resulting from any ideas, methods, instructions or products referred to in the content.

