## Supplementary Figures and Tables for "Integrated Multi-Omics Analysis for Molecular Subtyping in NSCLC: A Cohort Study": Suplementary materials.docx

**Table S1.** Subtype-specific mutated genes identified by independent tests of subtypes and mutations

| Gene  (Mutated) | TMB | CS1 | CS2 | CS3 | CS4 | CS5 | pvalue | padj |
| --- | --- | --- | --- | --- | --- | --- | --- | --- |
| *TP53* | 362  (66%) | 102  (87.9%) | 104  (92.0%) | 69  (49.3%) | 59  (47.2%) | 28  (49.1%) | 1.12e-23 | 6.93e-21 |
| *TTN* | 325  (59%) | 91  (78.4%) | 88  (77.9%) | 62  (44.3%) | 56  (44.8%) | 28  (49.1%) | 1.03e-12 | 2.13e-10 |
| *SYNE1* | 124  (22%) | 41  (35.3%) | 41  (36.3%) | 12  (8.6%) | 26  (20.8%) | 4  (7.0%) | 4.48e-10 | 6.93e-08 |
| *FAM135B* | 116  (21%) | 44  (37.9%) | 29  (25.7%) | 13  (9.3%) | 16  (12.8%) | 14  (24.6%) | 7.88e-08 | 8.13e-06 |
| *KRAS* | 86  (16%) | 1  (0.9%) | 0  (0.0%) | 27  (19.3%) | 43  (34.4%) | 15  (26.3%) | 1.37e-20 | 4.24e-18 |
| *KMT2D* | 84  (15%) | 28  (24.1%) | 28  (24.8%) | 9  (6.4%) | 16  (12.8%) | 3  (5.3%) | 7.00e-06 | 3.94e-04 |
| *KEAP1* | 81  (15%) | 17  (14.7%) | 6  (5.3%) | 18  (12.9%) | 19  (15.2%) | 21  (36.8%) | 1.45e-05 | 7.48e-04 |
| *NPAP1* | 77  (14%) | 19  (16.4%) | 4  (3.5%) | 23  (16.4%) | 17  (13.6%) | 14  (24.6%) | 6.26e-04 | 2.42e-02 |
| *ASXL3* | 65  (12%) | 8  (6.9%) | 5  (4.4%) | 20  (14.3%) | 18  (14.4%) | 14  (24.6%) | 6.88e-04 | 2.51e-02 |
| *LRRK2* | 56  (10%) | 15  (12.9%) | 26  (23.0%) | 8  (5.7%) | 4  (3.2%) | 3  (5.3%) | 3.85e-06 | 2.38e-04 |
| *STK11* | 45  (8%) | 0  (0.0%) | 1  (0.9%) | 19  (13.6%) | 14  (11.2%) | 11  (19.3%) | 7.36e-09 | 9.11e-07 |
| *CDKN2A* | 45  (8%) | 18  (15.5%) | 19  (16.8%) | 2  (1.4%) | 3  (2.4%) | 3  (5.3%) | 2.16e-07 | 1.83e-05 |
| *NFE2L2* | 44  (8%) | 17  (14.7%) | 19  (16.8%) | 1  (0.7%) | 5  (4.0%) | 2  (3.5%) | 2.36e-07 | 1.83e-05 |
| *EGFR* | 42  (8%) | 2  (1.7%) | 3  (2.7%) | 15  (10.7%) | 17  (13.6%) | 5  (8.8%) | 5.17e-04 | 2.13e-02 |
| *PLCH1* | 38  (7%) | 13  (11.2%) | 13  (11.5%) | 2  (1.4%) | 9  (7.2%) | 1  (1.8%) | 1.26e-03 | 4.33e-02 |
| *LPPR4* | 34  (6%) | 1  (0.9%) | 3  (2.7%) | 14  (10.0%) | 7  (5.6%) | 9  (15.8%) | 2.68e-04 | 1.18e-02 |
| *PTEN* | 34  (6%) | 15  (12.9%) | 14  (12.4%) | 1  (0.7%) | 4  (3.2%) | 0  (0.0%) | 1.60e-06 | 1.10e-04 |
| *CLCN1* | 32  (6%) | 3  (2.6%) | 0  (0.0%) | 12  (8.6%) | 9  (7.2%) | 8  (14.0%) | 1.93e-04 | 9.19e-03 |

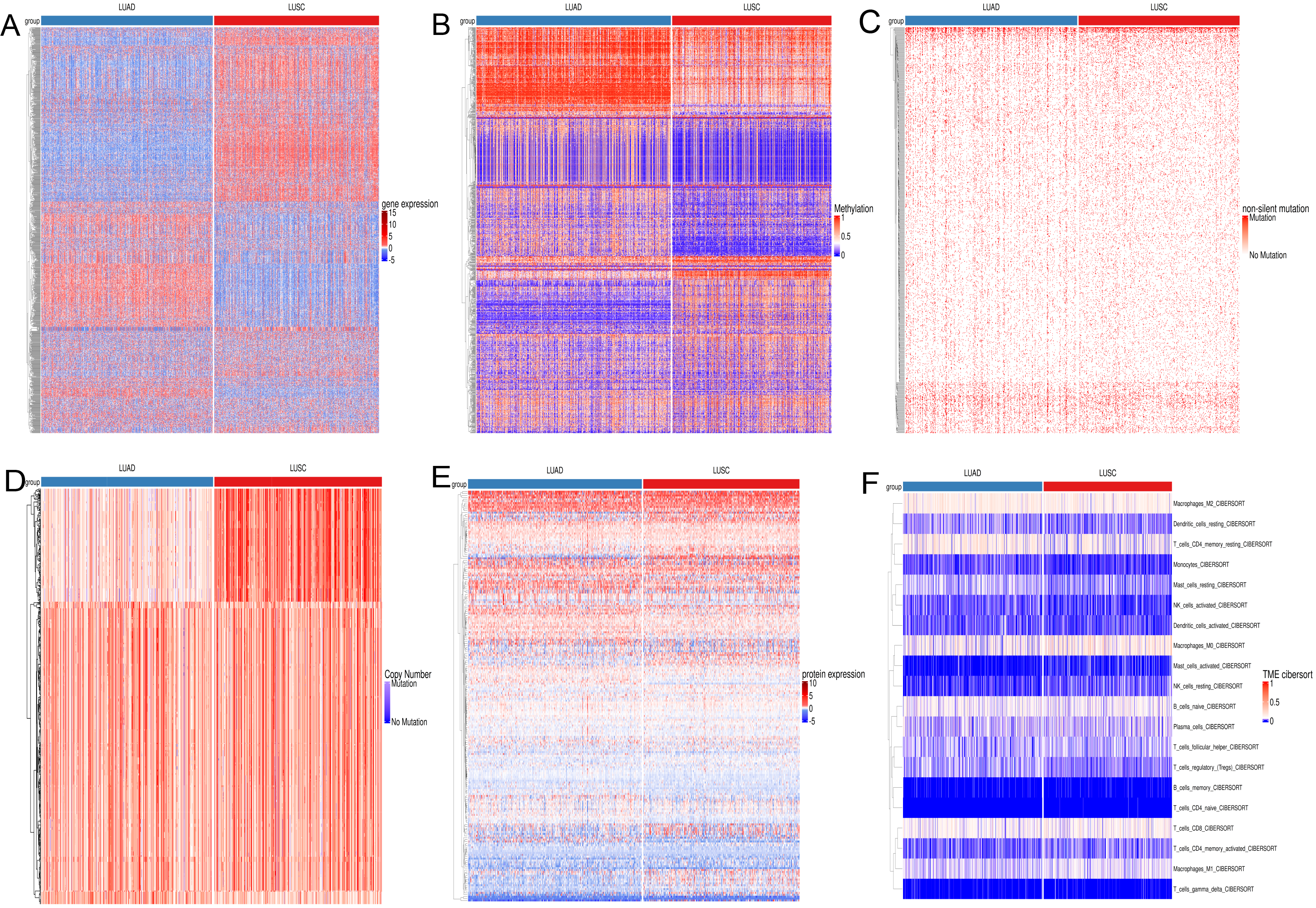

**Figure S1.** Heatmaps of six omics layers depicting the molecular landscapes distinguishing LUAD from LUSC. (**A**) heatmap of the top 1,000 highly variable genes based on gene expression data; (**B**) heatmap of β values for the top 1,000 CpG loci in DNA methylation data; (**C**) mutation heatmap of 651 genes mutated in at least 5% of samples in the somatic mutation data, with 0 indicating no mutation and 1 indicating presence of mutation; (**D**) copy number heatmap of the top 1,000 highly variable regions in CNV data; (**E**) heatmap of RPPA values for 216 proteins in the proteomics data; (**F**) heatmap of infiltration proportions of 22 immune cell types in the TME data, generated using the CIBERSORT algorithm.

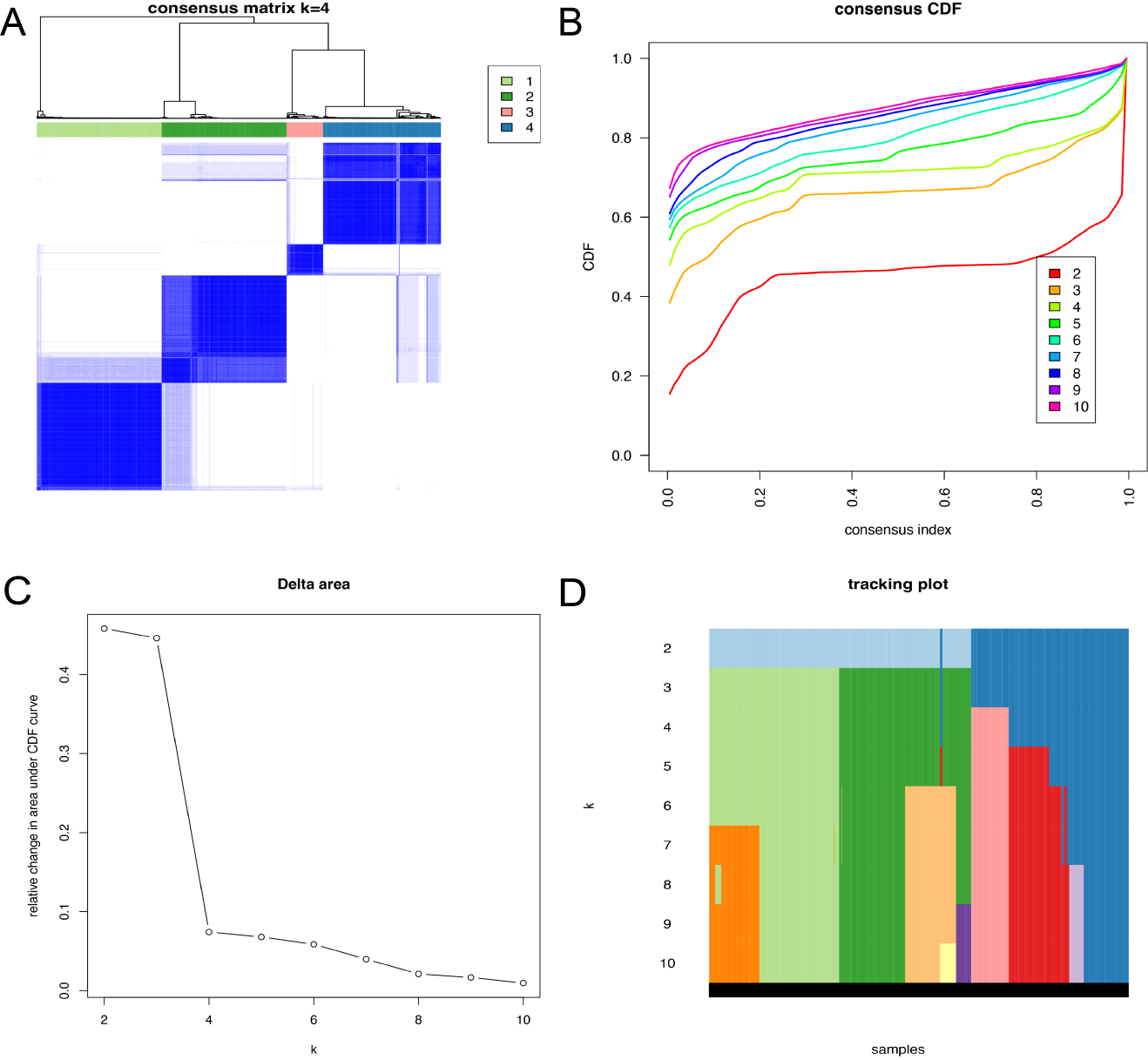

**Figure S2.** Consensus clustering results of DNA methylation data. (**A**) Consensus matrix heatmap at K = 4; (**B**) Cumulative distribution function plot; (**C**) Relative change in the area under the CDF curve (Delta Area Plot); (**D**) Sample classification tracking plot across different values of K.

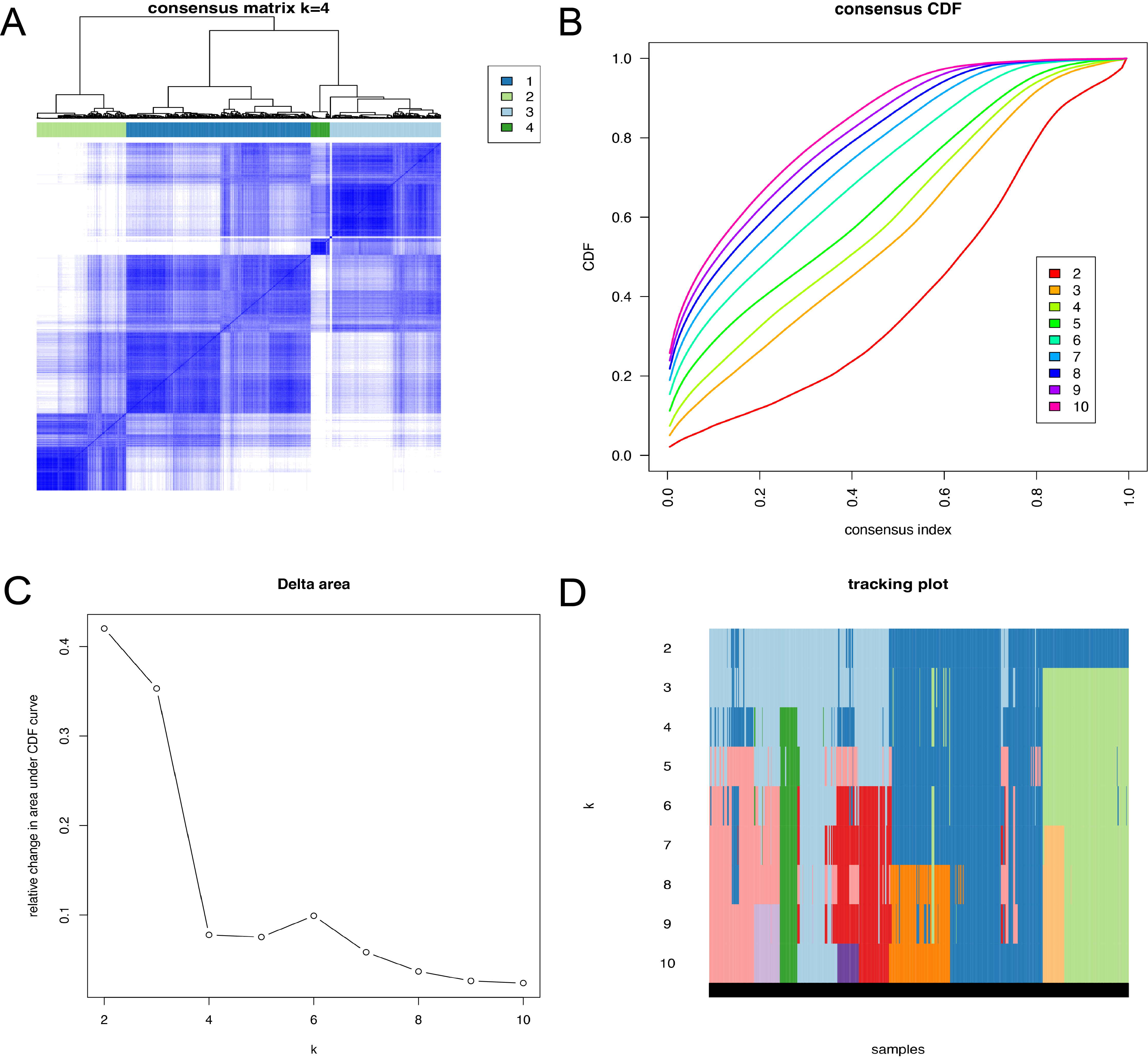

**Figure S3.** Consensus clustering results of somatic mutation data. (**A**) Consensus matrix heatmap at K = 4; (**B**) Cumulative distribution function plot; (**C**) Relative change in the area under the CDF curve (Delta Area Plot); (**D**) Sample classification tracking plot across different values of K.

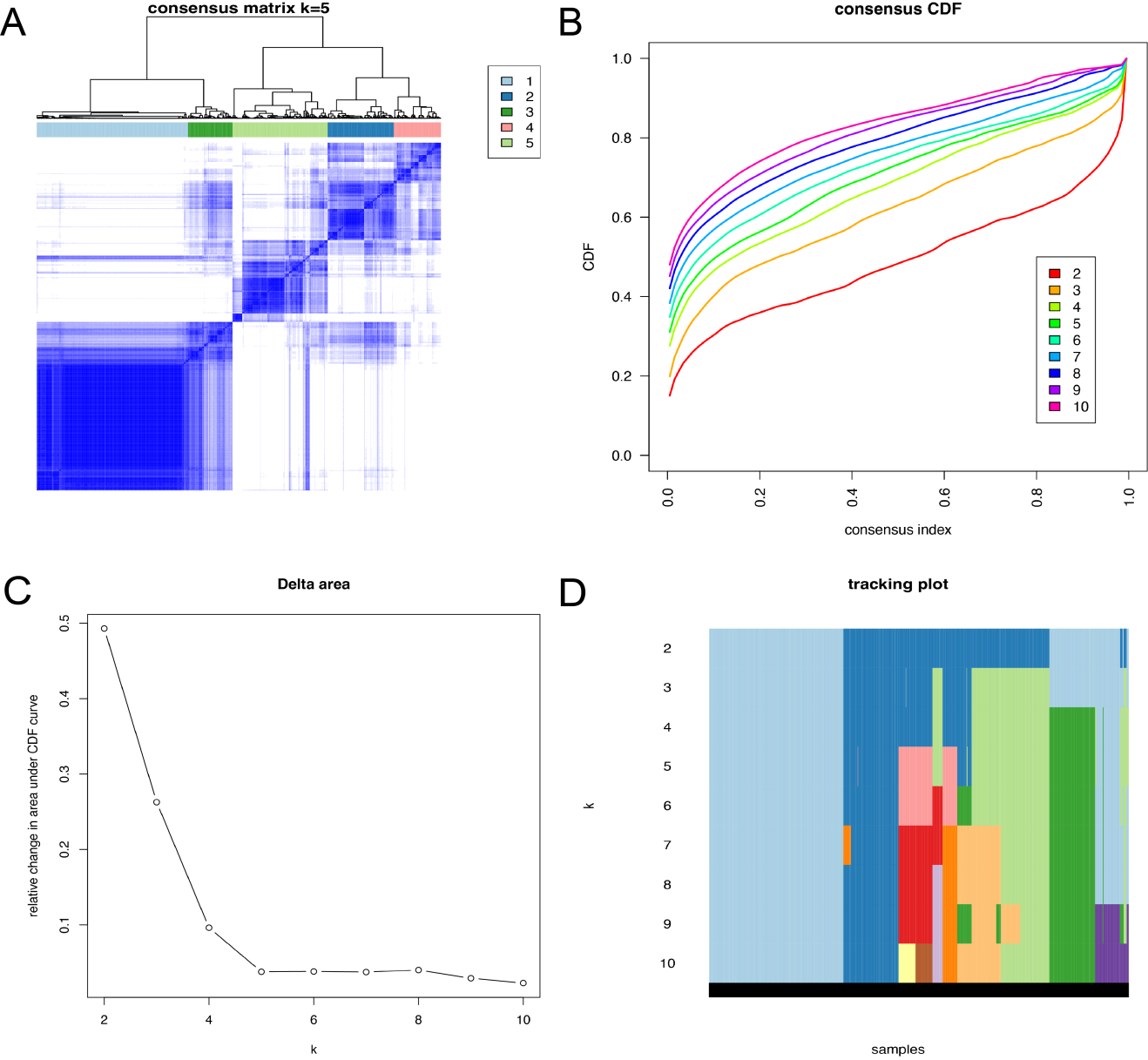

**Figure S4.** Consensus clustering results of CNV data. **(A**) Consensus matrix heatmap at K = 4; (**B**) Cumulative distribution function plot; (**C**) Relative change in the area under the CDF curve (Delta Area Plot); (**D**) Sample classification tracking plot across different values of K.

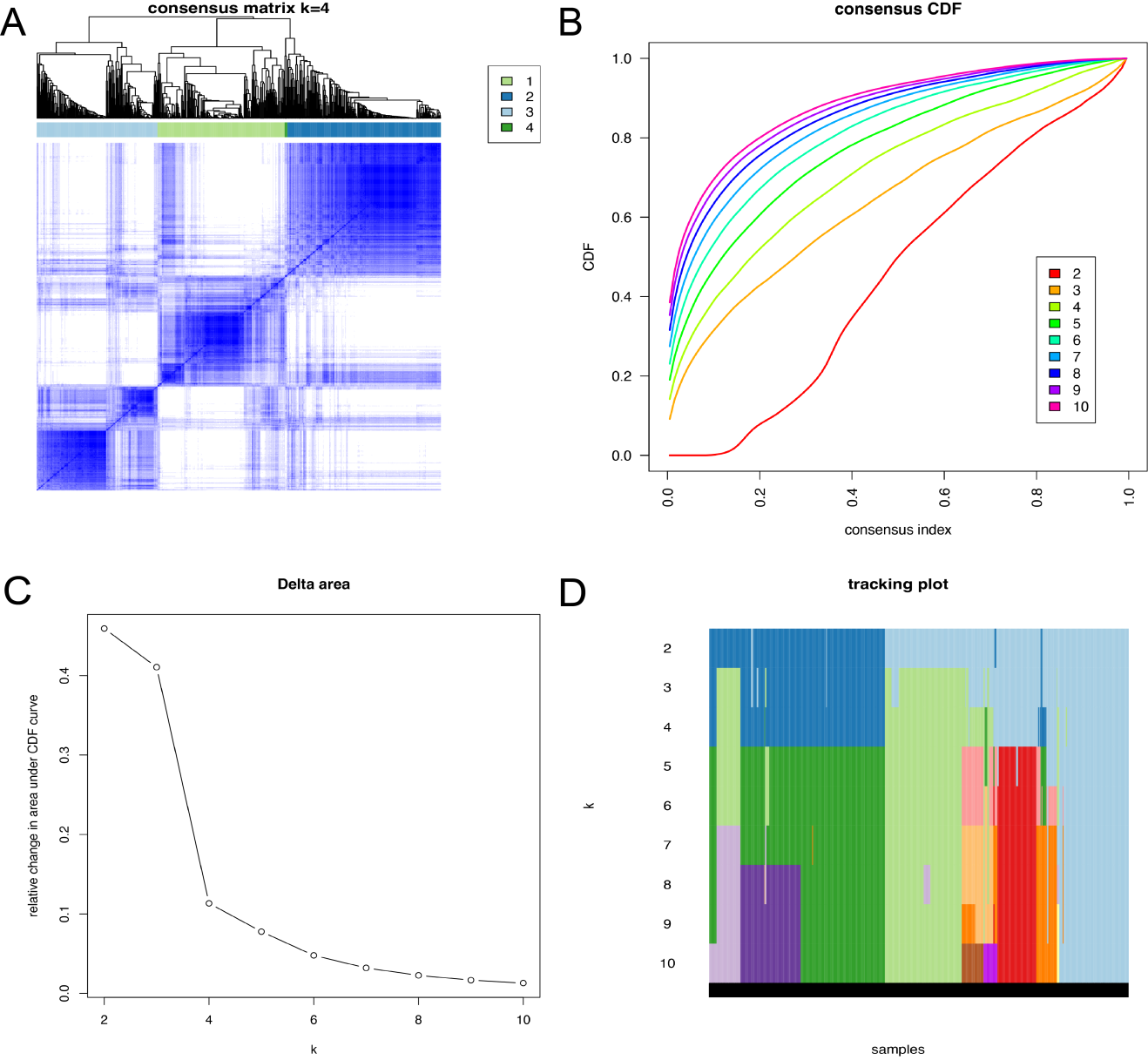

**Figure S5.** Consensus clustering results of protein expression data. (**A**) Consensus matrix heatmap at K = 4; (**B**) Cumulative distribution function plot; (**C**) Relative change in the area under the CDF curve (Delta Area Plot); (**D**) Sample classification tracking plot across different values of K.

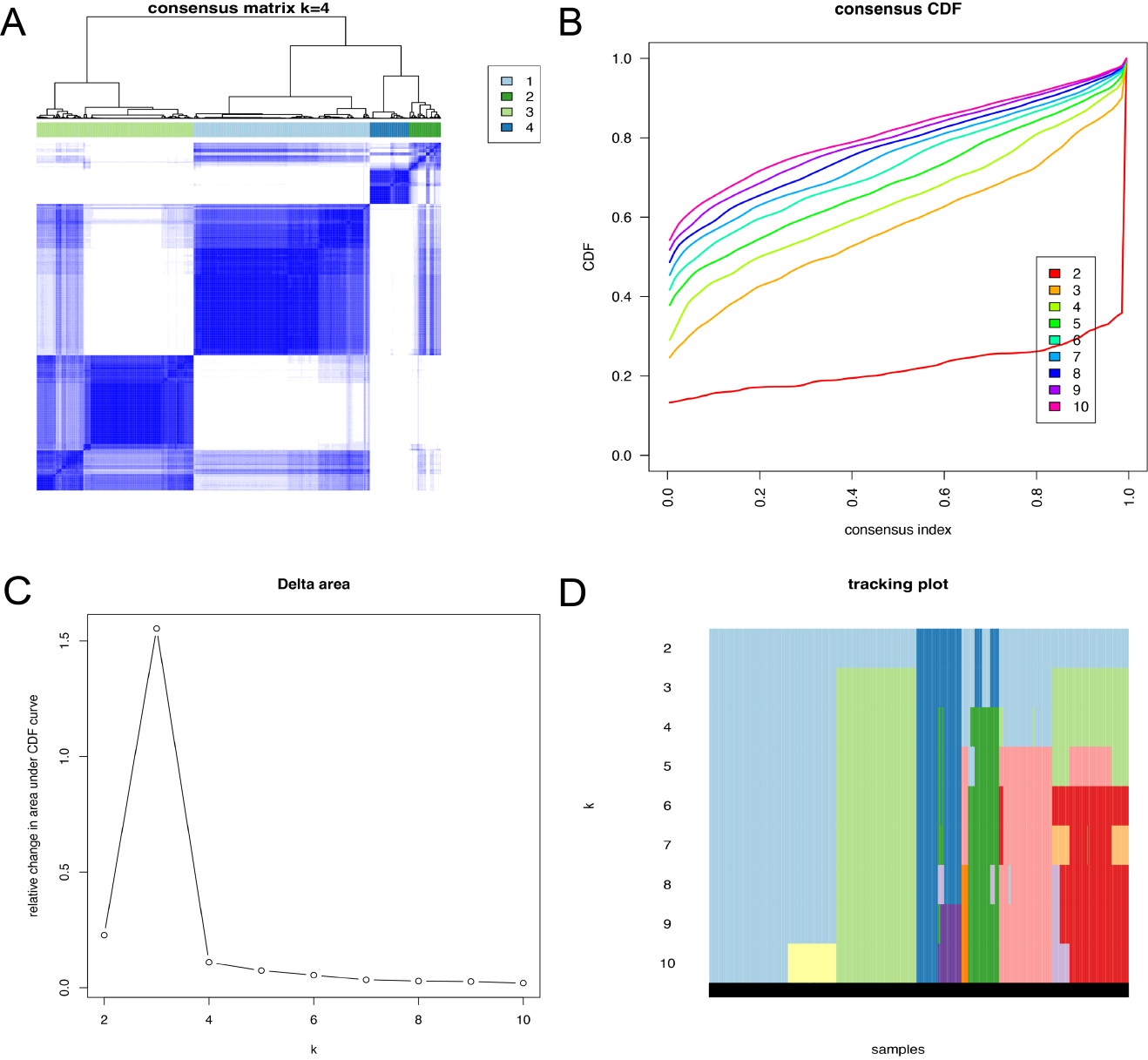

**Figure S6.** Consensus clustering results of TME data. (**A**) Consensus matrix heatmap at K = 4; (**B**) Cumulative distribution function plot; (**C**) Relative change in the area under the CDF curve (Delta Area Plot); (**D**) Sample classification tracking plot across different values of K.

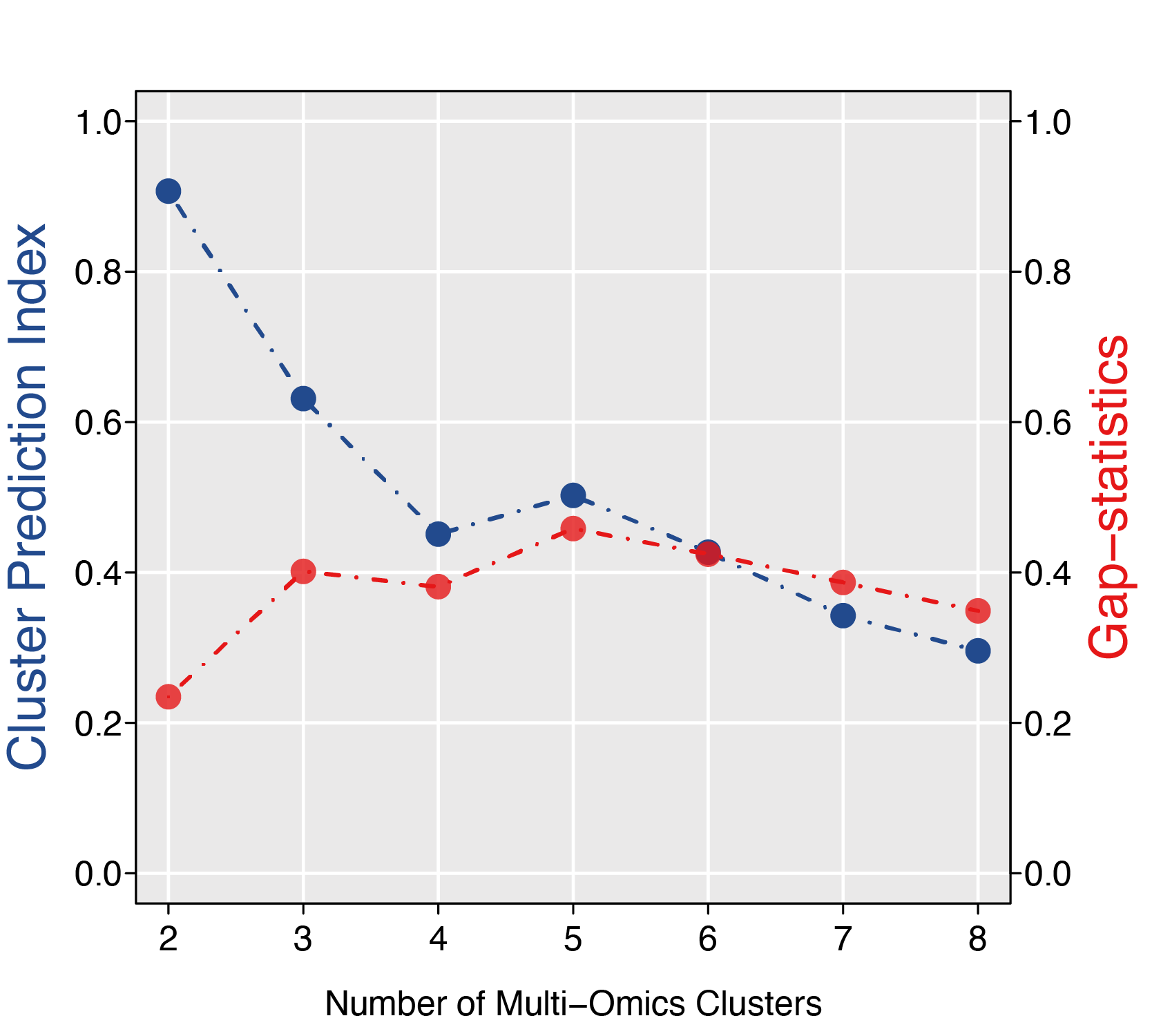

**Figure S7.** Line plots of CPI index and Gap statistic for multi-omics integrative clustering.The blue curve represents CPI values across different K values, while the red curve denotes Gap statistic values across different K values.

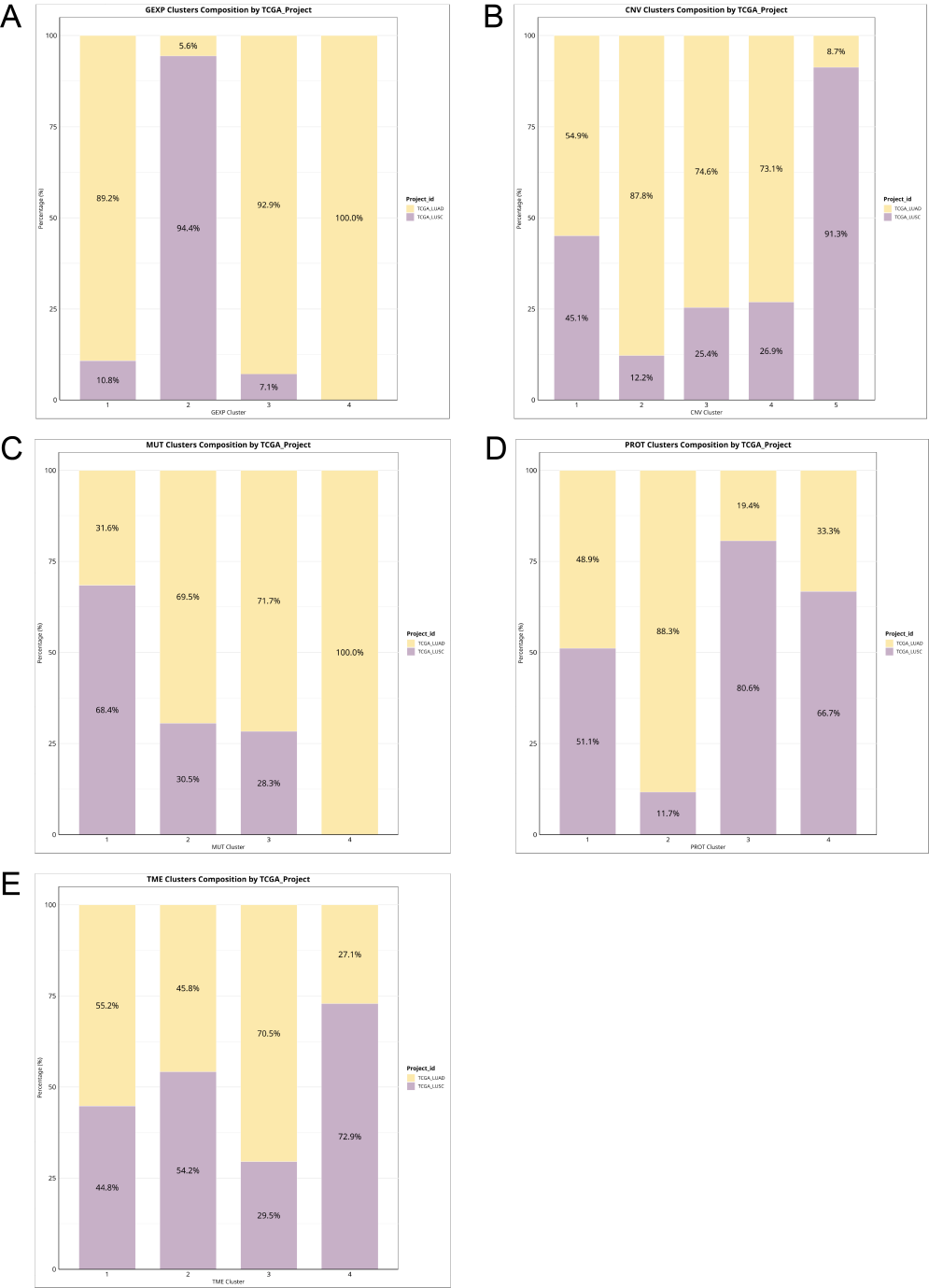

**Figure S8.** Distribution of histological subtypes (LUAD vs. LUSC) across the five single-omics consensus subtypes (excluding DNA methylation). (**A**) GEXP clusters 1 and 3 were enriched in LUAD, cluster 2 in LUSC, and cluster 4 contained only two LUAD cases. (**B**) Most CNV clusters (2, 3, 4) were LUAD-dominant, cluster 5 was LUSC-dominant, while cluster 1 showed a balanced distribution (LUAD 54.9%, LUSC 45.1%). (**C**) MUT clusters 2 and 3 were LUAD-dominant, whereas cluster 1 was LUSC-dominant; cluster 4 included only 30 LUAD cases. (**D**) PROT clusters 2 and 3 were dominated by LUAD and LUSC, respectively, while cluster 1 was relatively balanced, and cluster 4 contained only three samples. (**E**) For TME, clusters 1 and 2 were balanced, cluster 3 was LUAD-dominant (70.5%), and cluster 4 was LUSC-dominant (72.5%).

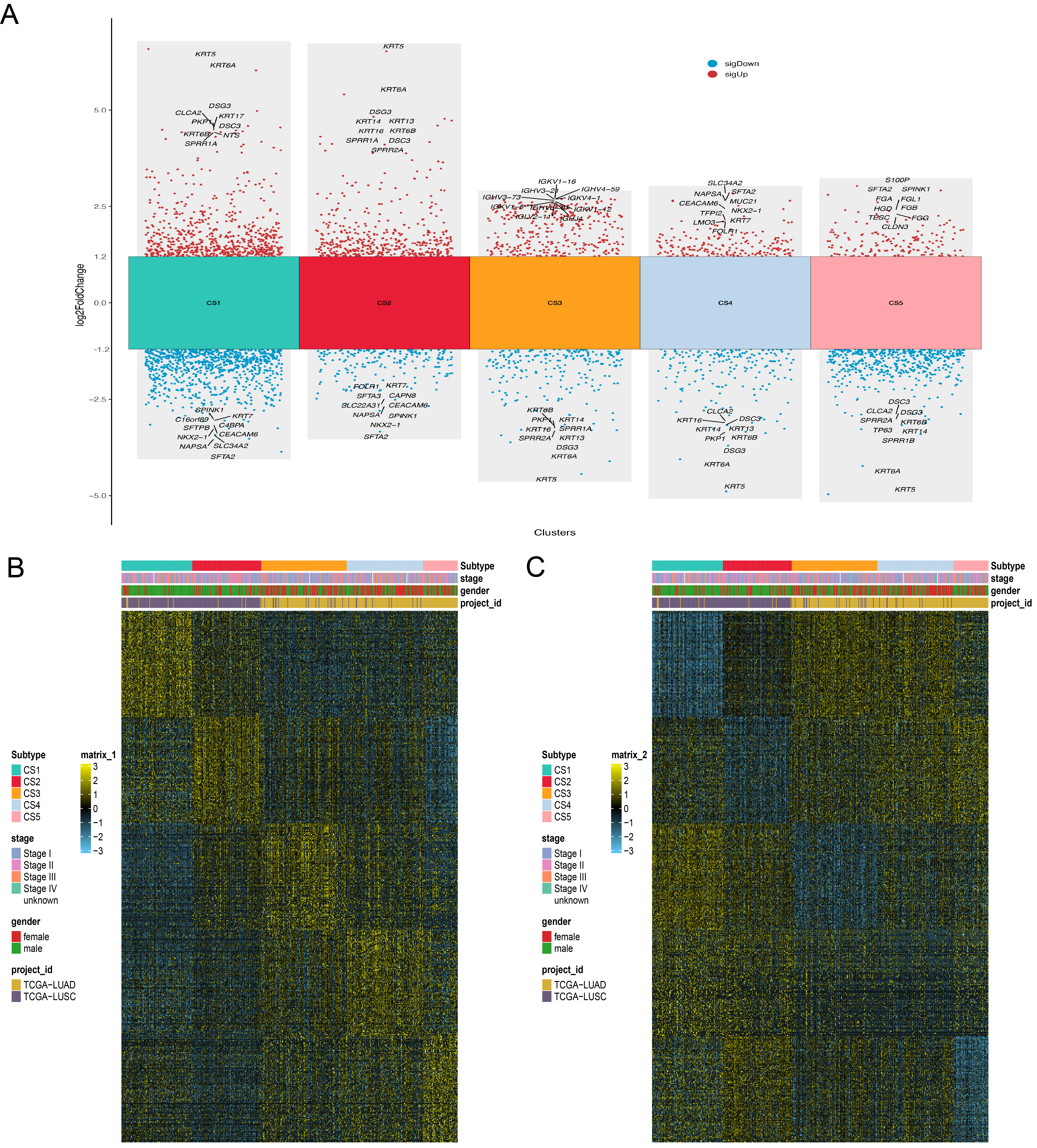

**Figure S9.** Identification of subtype-specific signature genes based on differentially expressed genes in each subtype. (**A**) Volcano plots of differentially expressed genes for the five multi-omics consensus subtypes. Differential genes were defined as |log2FoldChange| > 1.2 and p < 0.05; red dots indicate genes significantly upregulated in the current subtype, and blue dots indicate genes significantly downregulated. Non-significant genes are not shown. (**B**) Heatmap of the top 100 subtype-specific upregulated signature genes; (**C**) Heatmap of the top 100 subtype-specific downregulated signature genes. The criteria for selecting subtype-specific signature genes were the top 100 genes with the most significant log2FoldChange and non-overlapping with other subtypes.

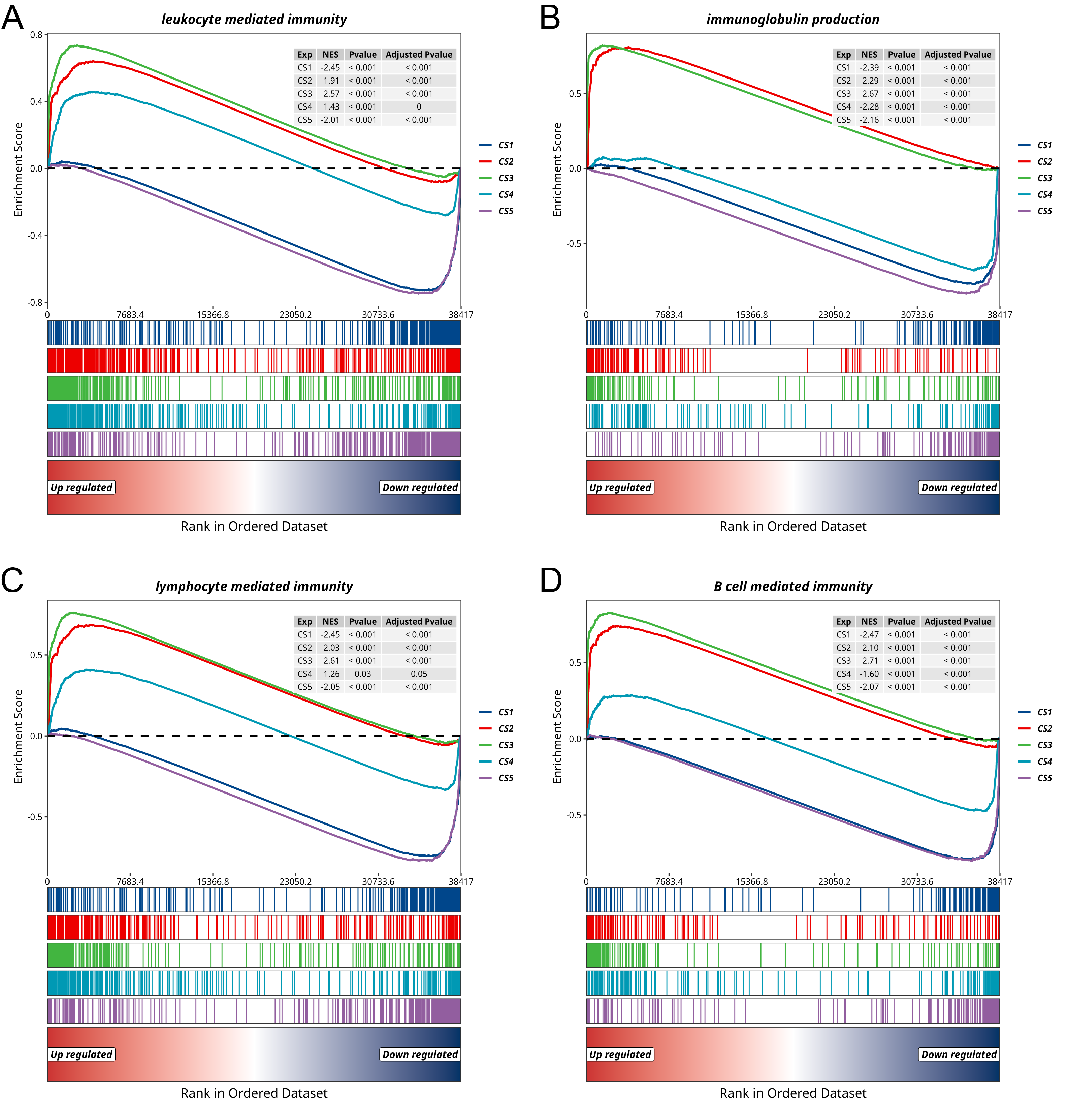

**Figure S10.** Comparison of selected immune-related GO terms enrichment among five multi-omics consensus subtypes. (**A**) Enrichment of the BP term “leukocyte-mediated immunity”; (**B**) Enrichment of the BP term “immunoglobulin production”; (**C**) Enrichment of the BP term “lymphocyte-mediated immunity”; (**D**) Enrichment of the BP term “B cell-mediated immunity.”
