## Supplementary figures and images for "Integrated Multi-Omics Analysis for Molecular Subtyping in NSCLC: A Cohort Study"

### Figure_S1.png

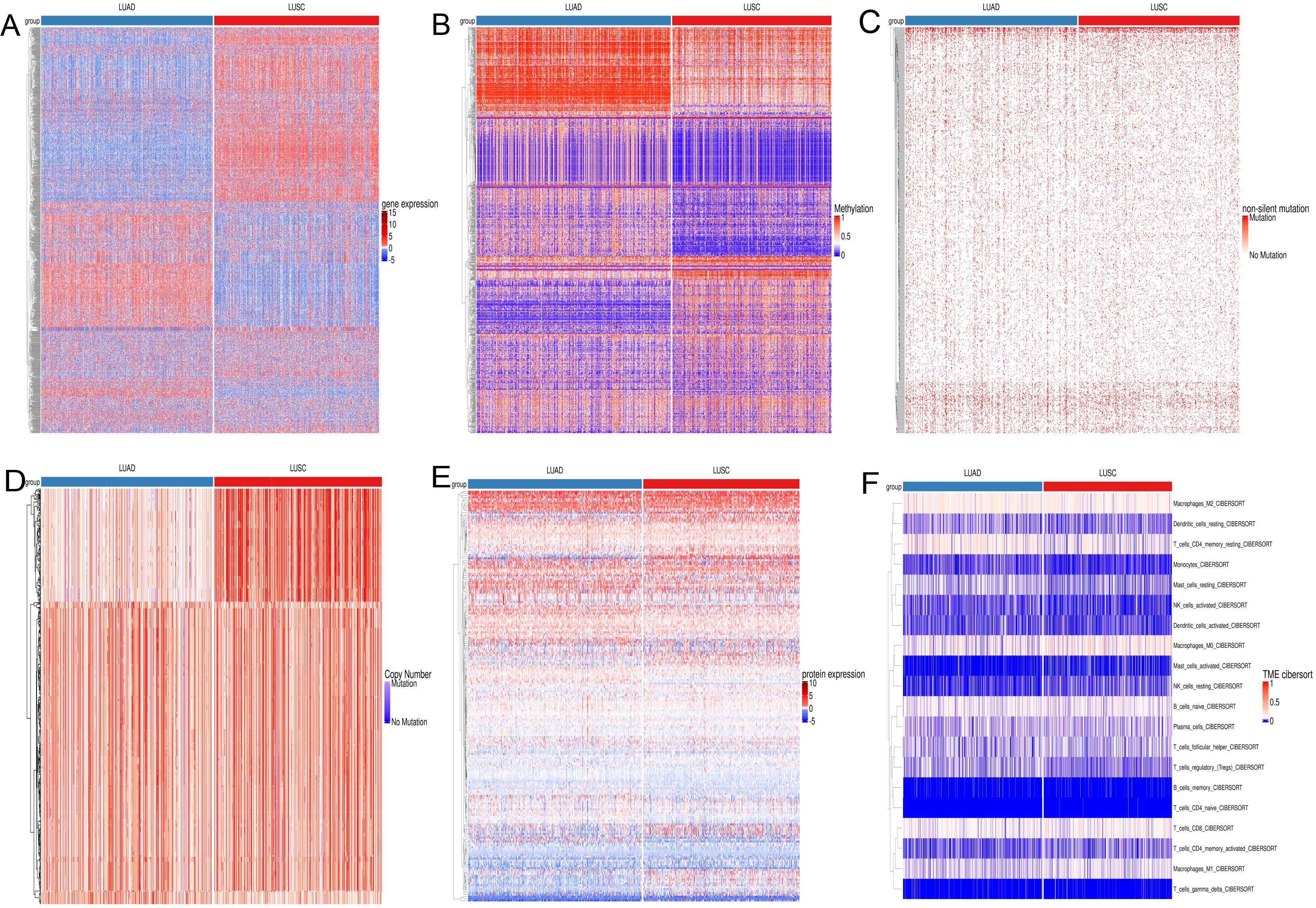

### Figure_S2.png

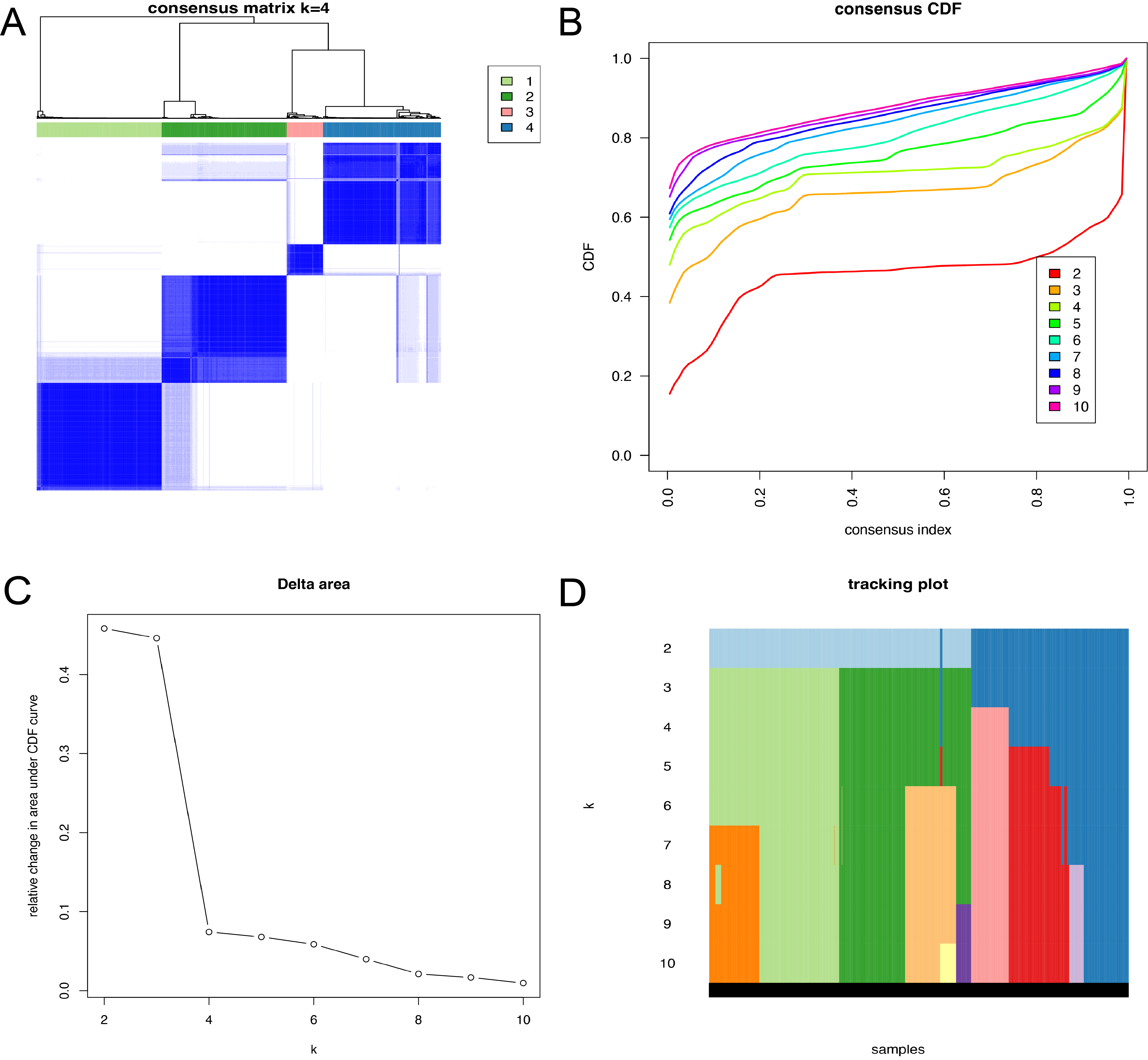

### Figure_S3.png

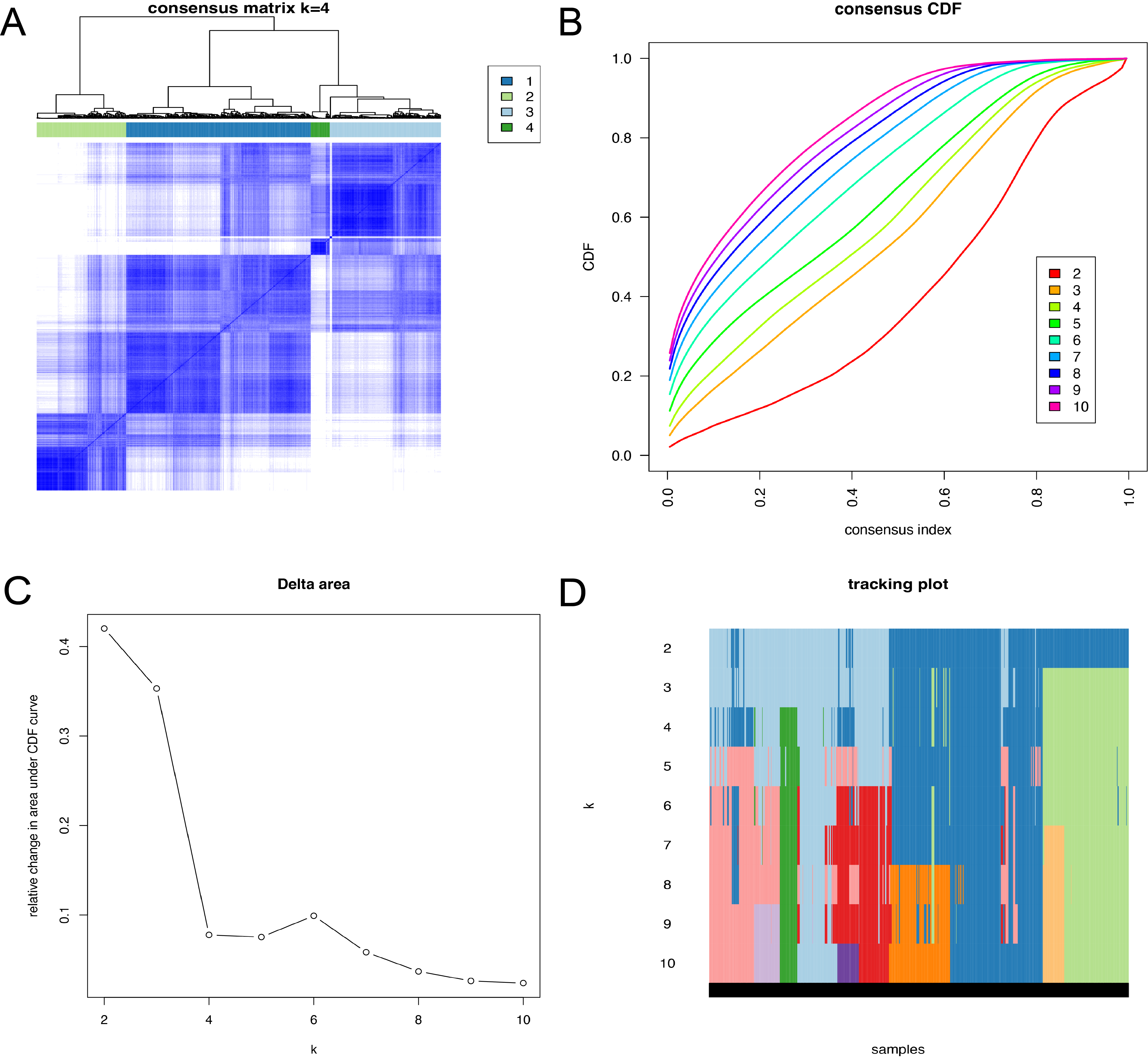

### Figure_S4.png

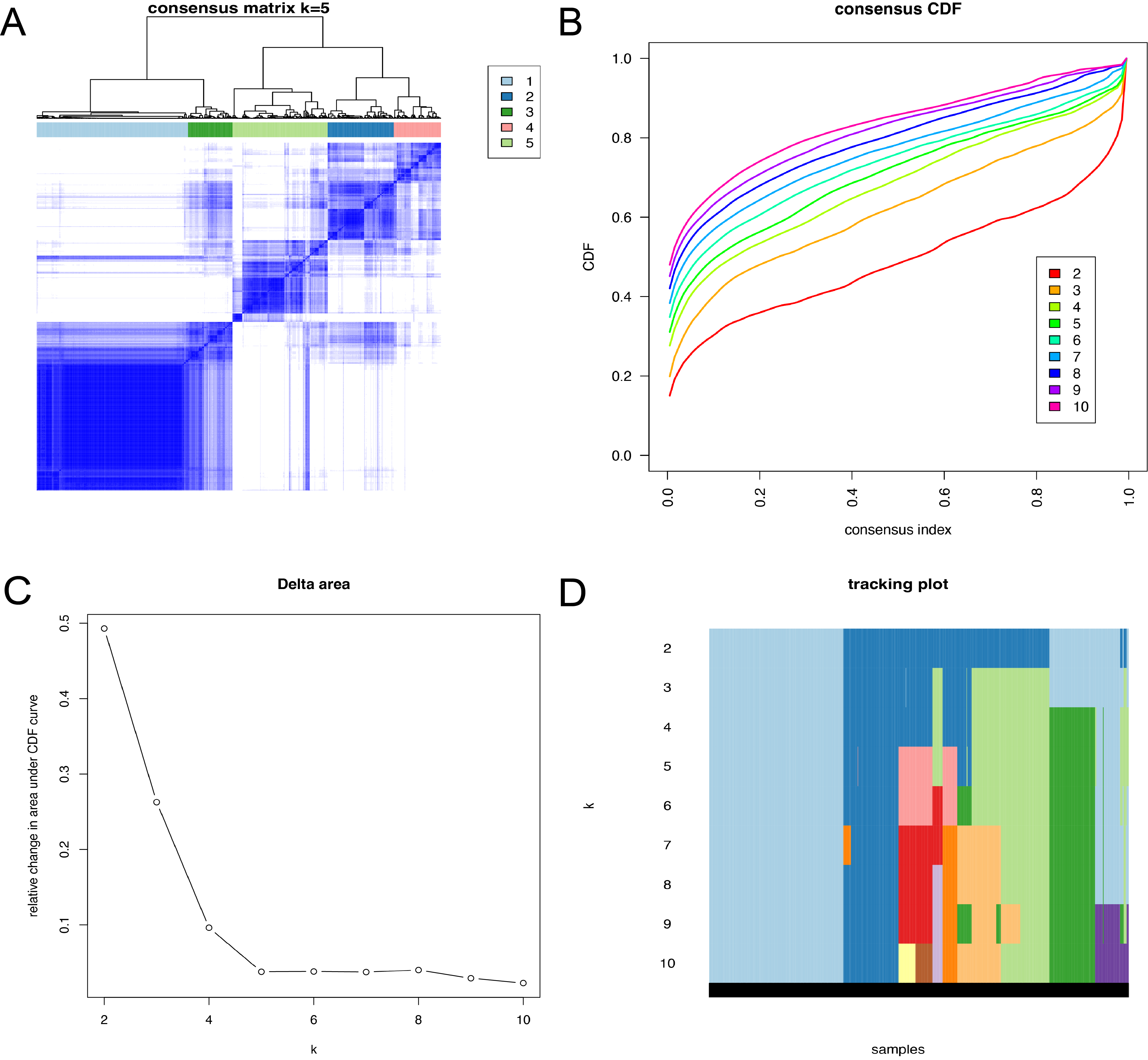

### Figure_S5.png

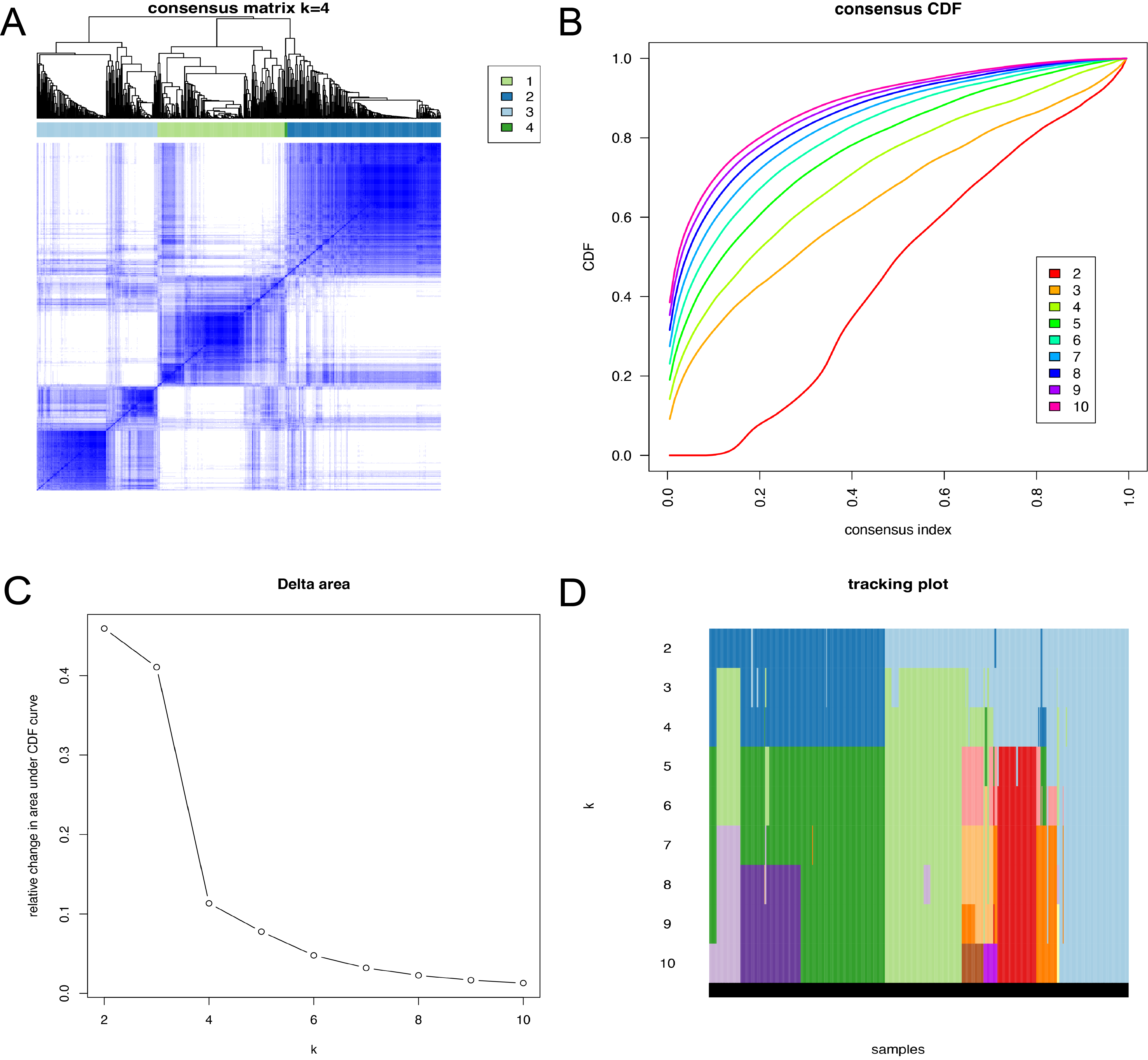

### Figure_S6.png

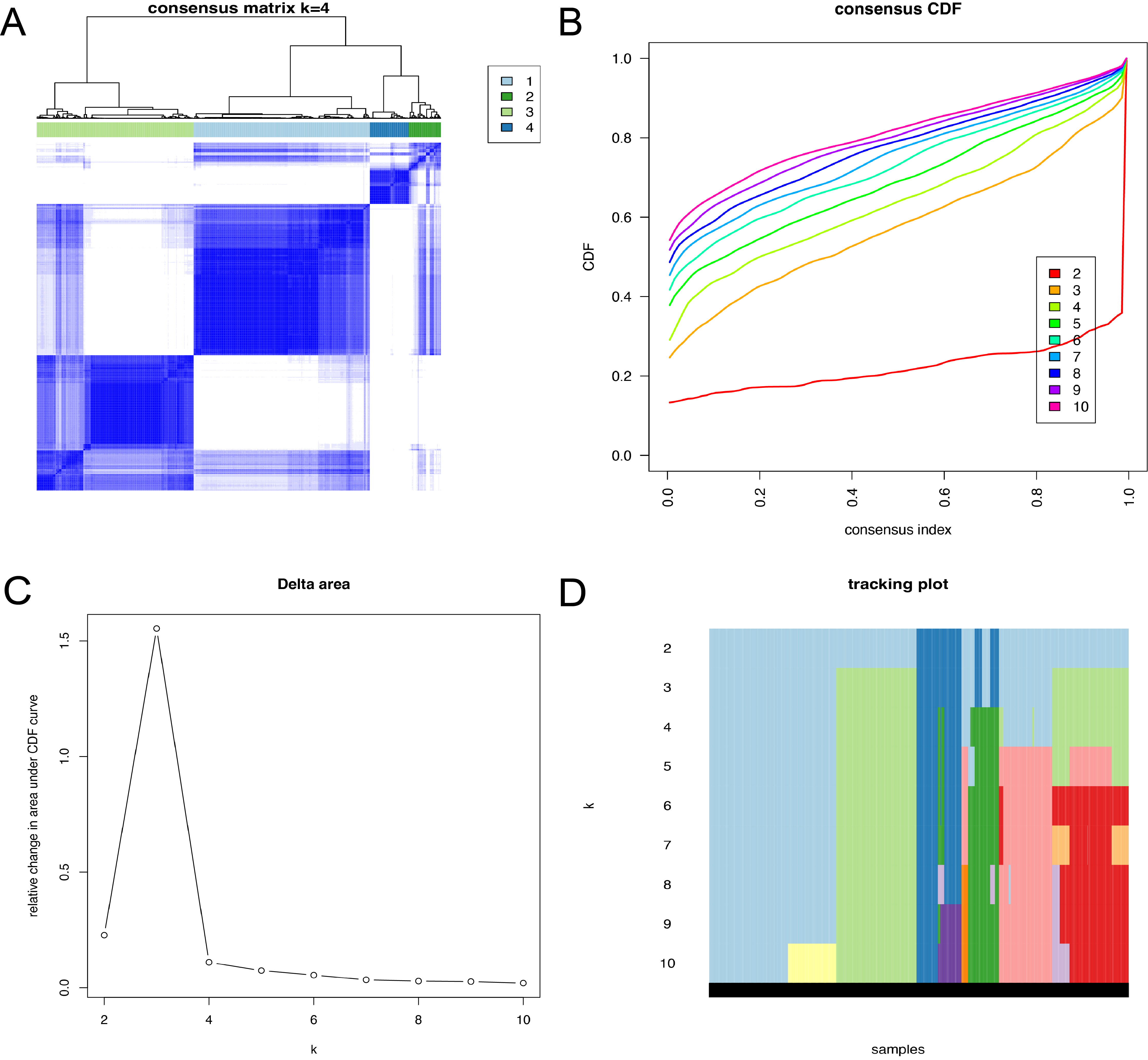

### Figure_S7.png

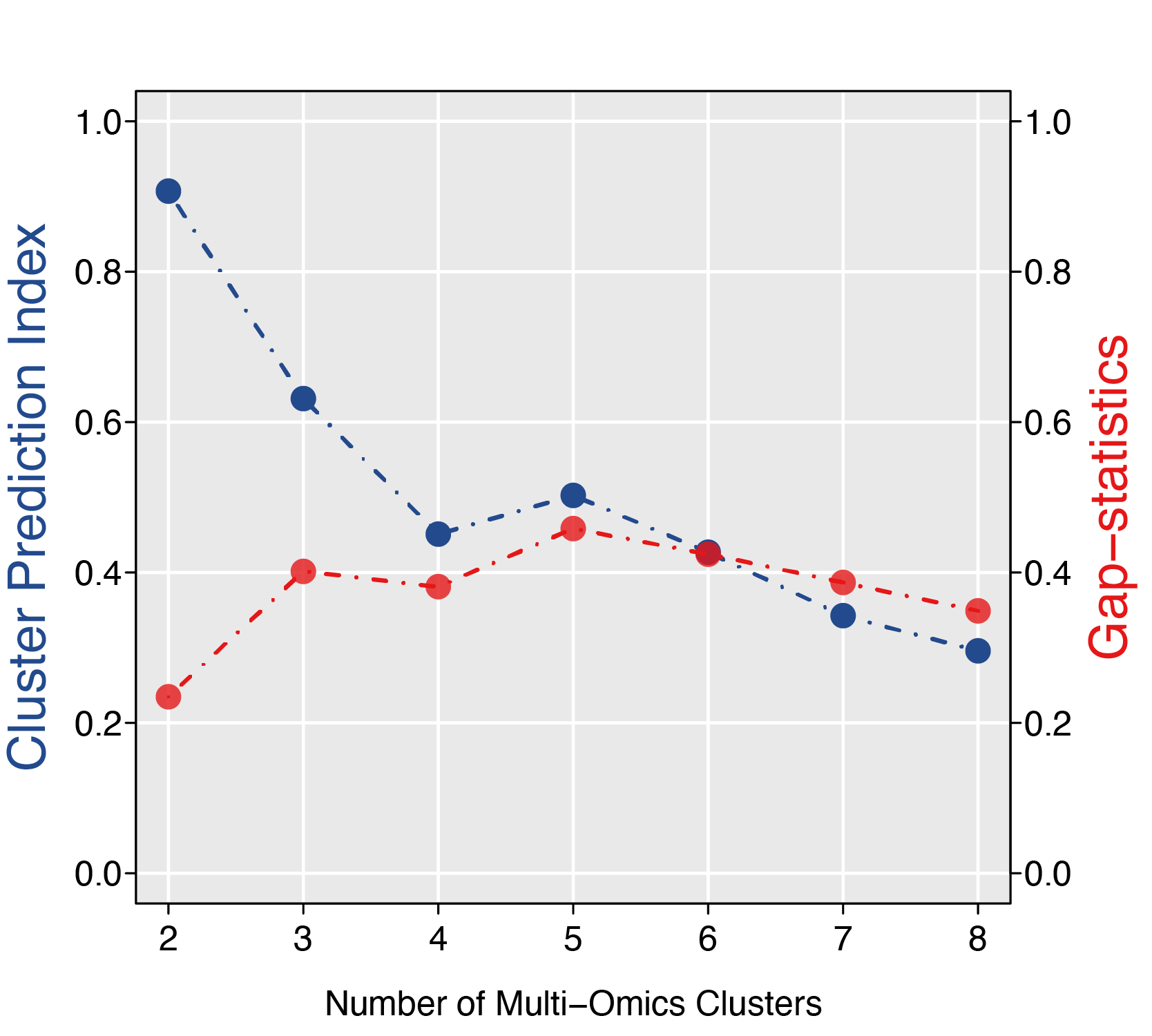

### Figure_S8.png

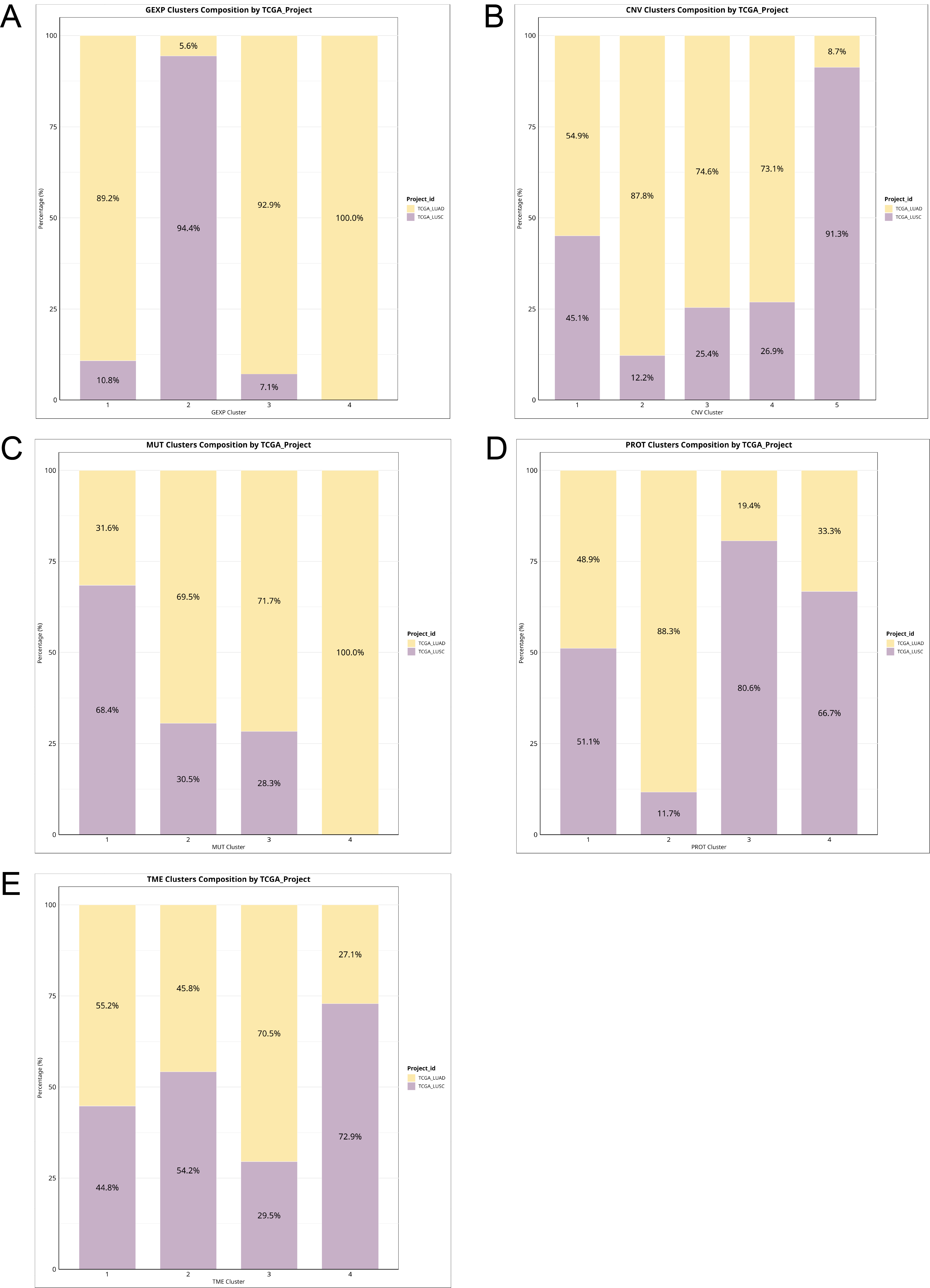

### Figure_S9.png

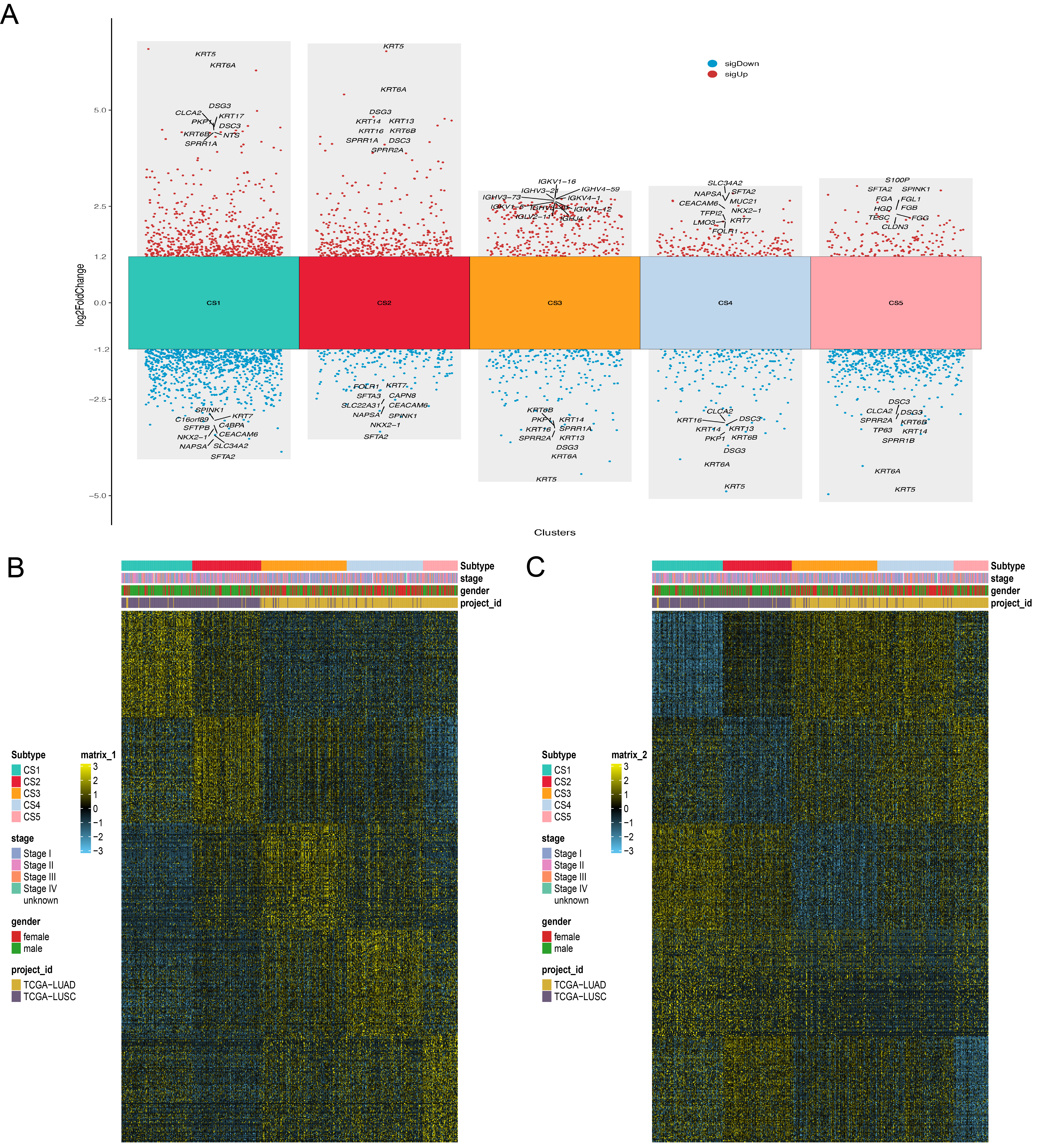

### Figure_S10.png

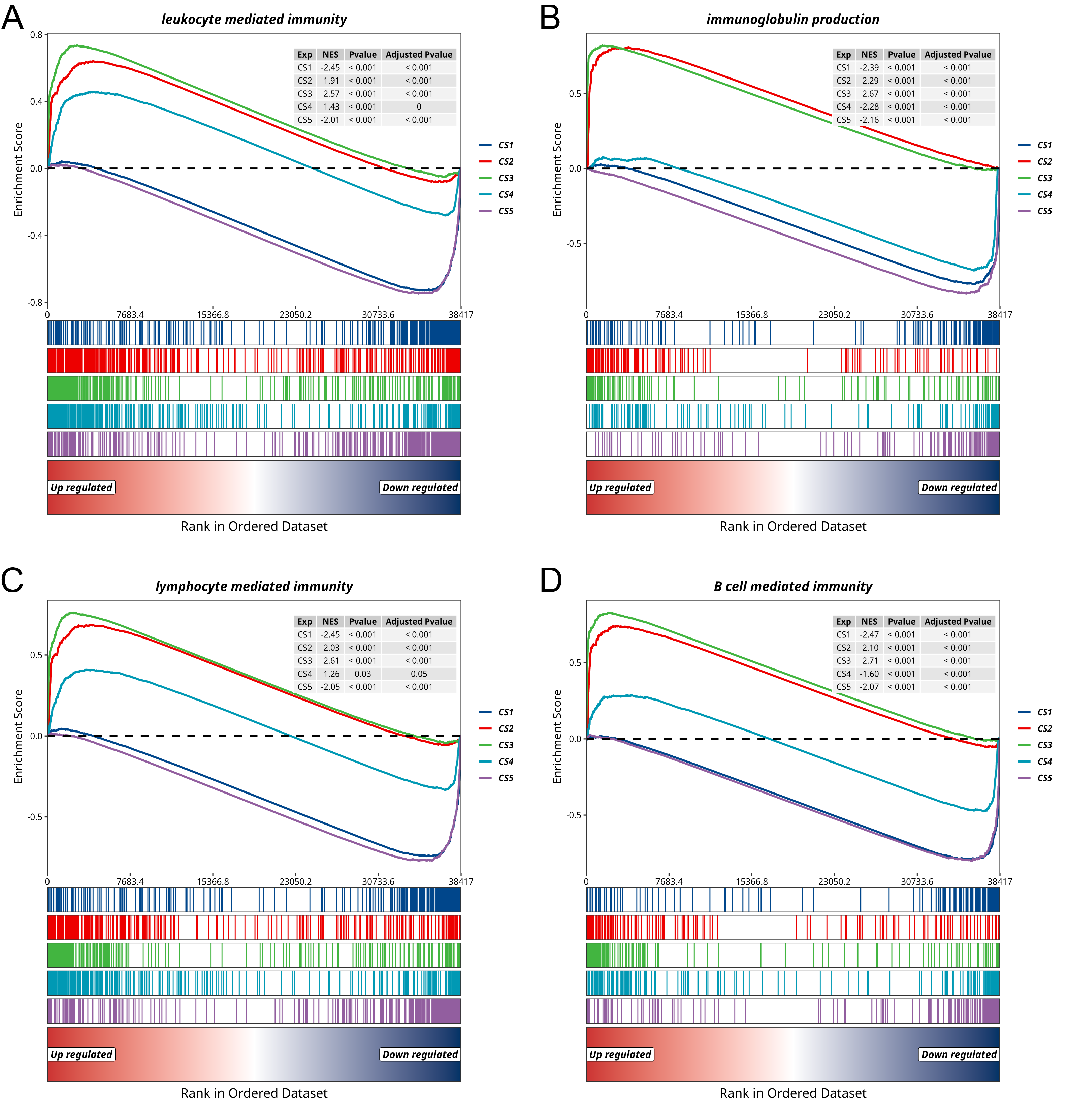
